# Inbreeding and demographic history of caribou (*Rangifer tarandus*) in western Canada inferred from genome-wide SNP data

**DOI:** 10.64898/2026.03.02.709119

**Authors:** Charlotte Bourbon, Samuel Deakin, Anita Michalak, Margaret M. Hughes, Maria Cavedon, Lalenia Neufeld, Agnès Pelletier, Jean Polfus, Helen Schwantje, Caeley Thacker, Marco Musiani, Jocelyn Poissant

**Author notes:** Corresponding authors: Charlotte Bourbon Email address, Jocelyn Poissant Email address, Marco Musiani.

## Abstract

Assessing genetic diversity is essential for conserving endangered populations, yet comprehensive genomic evaluations remain limited for many declining species. Here, we investigated inbreeding levels and effective population sizes (N_e_) of caribou (*Rangifer tarandus*) in western Canada, where populations have experienced pronounced declines over the past centuries due to anthropogenic pressures and climate change. We analyzed 33,346 Single Nucleotide Polymorphisms (SNPs) from 759 individuals representing 45 subpopulations within six metapopulations to: (1) assess inbreeding using runs of homozygosity (ROHs), (2) estimate contemporary and historical N_e_, and (3) evaluate relationships between census size (N_c_), inbreeding, and N_e_. Small and endangered subpopulations, predominantly in southern regions, generally exhibited high inbreeding (F_ROH_ > 0.1), although some larger populations also showed elevated levels. Most subpopulations displayed a mixture of short and long ROHs, indicating both ancient shared ancestry and recent inbreeding. Twelve subpopulations had N_e_ <50, and 28 subpopulations and all metapopulations had N_e_ < 500, suggesting compromised short-term viability and long-term adaptive potential. N_c_ significantly predicted inbreeding (*R*² = 0.25), whereas contemporary N_e_ did not. Historical N_e_ reconstructions revealed a north-to-south gradient in bottleneck timing: northern populations declined in ∼1700-1780, central populations in ∼1780-1860, and southern populations in ∼1860-1940, likely driven by sequential impacts of climate shifts and anthropogenic disturbances. Our findings identify at-risk populations requiring urgent genetic intervention and demonstrate that integrating inbreeding and N_e_ estimates provides a robust framework for caribou recovery and the management of fragmented wildlife populations.

## Introduction

Assessing genetic diversity is critical for conserving and managing at-risk populations, particularly as extinction rates accelerate globally (Allendorf et al., 2022). Genomic approaches have transformed wildlife conservation by enabling robust inference of key genetic predictors of both short- and long-term population persistence, particularly inbreeding and effective population size (N_e_, Hohenlohe et al., 2021). High-throughput single nucleotide polymorphism (SNP) genotyping offers advantages over traditional markers like microsatellites, providing greater genomic coverage and higher resolution for estimating inbreeding coefficients and N_e_ (Forneris et al., 2025). This improved accuracy facilitates better detection of associated risks, such as inbreeding depression and the loss of genetic diversity through drift, which are critical concerns for endangered species conservation (Kardos et al., 2021). Integrating genomic data with demographic information also allows for reconstructing population histories and identifying critical thresholds (Chen et al., 2025), ultimately informing adaptive management strategies to preserve genetic diversity and promote population resilience (Hoffmann et al., 2015; Hohenlohe et al., 2021).

Among the genetic threats facing at-risk populations, inbreeding poses a particularly severe risk to population viability (Forneris et al., 2025). By reducing genetic diversity and fitness-associated overdominance, while increasing the expression of deleterious recessive alleles, inbreeding can undermine the survival of small populations (Charlesworth & Willis, 2009). One method useful for detecting inbreeding is the analysis of runs of homozygosity (ROHs). ROHs are continuous genomic segments where homologous chromosomes are identical due to inheritance from a common ancestor (Curik et al., 2014). They provide detailed insights into both individual and population inbreeding levels in the absence of pedigree data (Kardos et al., 2015), as is usually the case with free-ranging wildlife species. Unlike pedigree analysis, which underestimates inbreeding due to incomplete records, and heterozygosity metrics like F_IS_, which only capture population-level averages, ROH analysis provides individual-level estimates (Shafer & Kardos, 2025). ROH are shaped by recombination and historical demographic events, and their length distributions reflect underlying coalescent history: long ROHs indicate recent inbreeding while short ROHs capture distant ancestry (Howrigan et al., 2011; Kumar et al., 2023). ROHs are widely used in livestock genetics (Peripolli et al., 2017) and have gained traction in wildlife conservation, including recent studies in red deer (*Cervus elaphus*; Hewett et al., 2024), feral horses (*Equus caballus*; Colpitts et al., 2022), and killer whales (*Orcinus orca*; Foote et al., 2021).

Beyond direct measures of inbreeding, N_e_ is another widely used metric for understanding genetic processes that shape population viability, such as genetic drift (Lande & Barrowclough, 1987; Waples, 2025). N_e_ is defined as the size of an ideal Wright-Fisher population experiencing the same rate of genetic drift or inbreeding as the population under study (Wright, 1931; Caballero, 2020). Unlike census population size (N_c_), N_e_ represents the number of breeding individuals contributing to the genetic makeup of a population (Caballero, 2020). In conservation genetics, N_e_ is often interpreted through the 50/500 rule, which suggests that N_e_ ≥50 prevents inbreeding depression over short timescales, while N_e_ ≥500 maintains long-term evolutionary potential (Frankham, 2014; Franklin, 1980). While recent debate suggests these thresholds may need revision (Waples, 2025), the 50/500 framework provides widely-used benchmarks for identifying at-risk populations and facilitates comparison among studies (Clarke et al., 2024). Importantly, this rule can also be applied to metapopulations, considering both migration and resulting gene flow (Kurland et al., 2023).

In addition to N_e_ itself, another important indicator is the N_e_/N_c_ ratio (Wright et al., 2021). This ratio is commonly used to assess the extent of genetic variation loss, as lower values indicate stronger effects of genetic drift over time (Mastretta-Yanes et al., 2024), and to provide a broader understanding of population dynamics (Luikart et al., 2010). For example, a population can have a low N_e_ despite a high N_c_ due to factors such as uneven sex ratios, variation in reproductive success, non-random mating, and overlapping generations (Waples 2024b). The N_e_/N_c_ ratio thus highlights populations in which demographic or ecological factors may negatively affect the maintenance of genetic diversity (Frankham, 1995; Jamieson & Allendorf, 2012). In wild populations, including both plants and animals, this ratio typically ranges between 0.1 and 0.2 (Frankham et al. 2014). When it falls below 0.1, populations may face a higher risk of inbreeding depression and genetic erosion, raising concerns for short- and long-term viability (Garner et al., 2020; Luikart et al., 2010).

While estimates of contemporary N_e_ have long been possible, genomic data such as medium- or high-density SNP genotypes now provide more precise estimates and enable reconstruction of historical trends in N_e_ (Nadachowska-Brzyska et al., 2022; Waples, 2024a). For instance, new methods combining LD and recombination rates can infer N_e_ changes over the last 100 generations with high accuracy (Quinn et al., 2024). Recent declines in N_e_ may be indicative of inbreeding and/or reduction in N_c_ (Robinson et al., 2019). These declines can sometimes be intensified by anthropogenic pressures (Crossman et al., 2024) or climatic events (Balza et al., 2025). N_e_ can also decline over time even when N_c_ remains stable, suggesting that genetic bottlenecks can occur without drastic changes in overall population numbers (Hoelzel, 1999; Hoelzel et al., 2024). For example, in human populations, founder effects and selective factors, including assortative mating and migration, can lead to reduced genetic diversity and N_e_ even when contemporary N_c_ are large (Browning et al., 2018). Short- and long-term conservation strategies can benefit from understanding historical N_e_ trends, as these can reveal population and genetic dynamics over time and prioritize interventions such as genetic rescue or conservation breeding to avoid population extinction (Allendorf et al., 2022). For example, a study on grizzly bears (*Ursus arctos*) found that N_e_ trends reflected the impact of management actions such as habitat restoration, whereas N_c_ failed to capture genetic recovery, aiding in the design of future conservation strategies (Kamath et al., 2015).

Caribou (*Rangifer tarandus*) are iconic ungulates native to North America and important to tundra, boreal and mountain ecosystems (Drever et al., 2019). In central British Columbia, caribou were relatively widespread until the late 1800s when their distribution began to decline, coinciding with Euro-American settlement expansion (Santomauro et al., 2012), a pattern observed across much of western Canada (Bergerud, 1974). Most populations have declined further over the past four decades, from approximately 40,000 to 15,000 individuals (Ministry of Forests, 2018), hypothesized to be due to anthropogenic and environmental stressors (Spalding 2000; Thomas and Gray 2002). In British Columbia, 11 subpopulations were classified as extirpated or functionally extirpated in 2023 (Government of British Columbia, 2025), while others are classified as being of special concern or threatened under the Canadian Species at Risk Act (Government of Canada, 2002). Habitat changes resulting from resource extraction activities and road infrastructure are the primary drivers of decline, disrupting movement and altering food availability (Hebblewhite 2017). These disturbances also increase vulnerability to wolf (*Canis lupus*) predation through improved predator access via anthropogenic linear features and apparent competition with other ungulates thriving in disturbed landscapes (Neufeld et al., 2021).

Caribou management actions in British Columbia shifted toward explicit conservation and recovery approaches in the late 1970s (Bergerud, 1974, 1978), with structured recovery plans for mountain and boreal caribou endorsed in the 2000s, followed by the province-wide Caribou Recovery Program in 2017 (Environment Canada, 2014; Ministry of Forests, 2018). In Alberta, Parks Canada implemented conservation measures in Jasper National Park since 2006, including access restrictions, predator pathway management, and disturbance reduction, addressing caribou declines partly driven by historical elk management policies and subsequent wolf population increase (Parks Canada, 2023). Concurrently, the Government of Alberta has undertaken habitat protection and predator management across provincial lands. Local conservation actions, including predator control in British Columbia and Alberta (B.C. Ministry of Water, Land and Resource Stewardship, 2024; Lamb et al., 2022), maternity penning (Lamb et al., 2022), or primary prey management (McLellan et al. 2025), can stabilize or boost subpopulations in the short-term. However, demographic improvements have not necessarily translated into improved genetic diversity, as genetic recovery typically lags due to limited gene flow and potential inbreeding in small populations (Robinson et al. 2023).

Understanding inbreeding and N_e_ patterns in caribou subpopulations and metapopulations is crucial for their conservation. In western Canada, while considerable research has focused on caribou population structure (Deakin et al., 2025; Taylor et al., 2021), genetic diversity (Taylor et al., 2024) or ecological factors such as habitat selection, migration, and behavior (Cavedon et al., 2022; Hughes et al., 2025; Theoret et al., 2022), studies on genetic indicators such as inbreeding and N_e_ remain limited (Cavedon et al., 2023; Taylor et al., 2024). Further investigations across additional subpopulations and metapopulations are needed, particularly regarding the genetic consequences of recent bottlenecks.

In this study, we used single nucleotide polymorphism (SNP) array data to assess contemporary and historical inbreeding and N_e_ across caribou subpopulations and metapopulations in western Canada. Our objectives were to: (1) identify subpopulations and metapopulations at risk based on inbreeding levels, contemporary N_e_, and the N_e_/N_c_ ratio; (2) investigate inbreeding and temporal changes in N_e_ over the past 100 generations to assess the genetic impact of key demographic events; and (3) assess the extent to which subpopulations and metapopulations share similar genetic and demographic histories. We hypothesized that smaller, more isolated subpopulations would exhibit higher inbreeding and lower N_e_ and predicted that this would be reflected in longer and more frequent ROHs in peripheral or fragmented subpopulations compared to larger, more connected ones. We further hypothesized that inbreeding and N_e_ are both driven by demographic change and predicted that temporal patterns in inbreeding and N_e_ would closely mirror fluctuations in N_c_ across subpopulations. Finally, we hypothesized that shared large-scale climatic and anthropogenic influences have generated comparable temporal trends in N_e_ among subpopulations and metapopulations across western Canada and predicted that N_e_ trajectories would show synchronous declines corresponding to known periods of habitat disruption and population fragmentation. Together, these analyses provide critical insights for guiding short- and long-term conservation strategies for caribou in western Canada.

## Materials and Methods

### Sample collection and genotyping

The samples analyzed here were previously described by Deakin et al. (2025). Briefly, blood or tissue samples were collected from 862 individuals across 45 caribou subpopulations in British Columbia and Alberta, Canada, by the Government of British Columbia, Parks Canada, and First Nations partners during free-ranging animal captures conducted between 2012 and 2024 (Figure 1). After quality control (see below), 759 individuals were retained for analysis. Samples included four extirpated subpopulations: Maligne was extirpated in 2020, Purcells South in 2021, South Selkirks in 2019, and Banff in 2009 (Government of British Columbia, 2025; Parks Canada, 2023). We selected subpopulations as our research unit, as they represent the local management units used by the Government of British Columbia (reviewed in Weckworth et al., 2018). The 45 subpopulations were also pooled into six metapopulations identified using hierarchical population structure analyses by Deakin et al. (2025): North-eastern, North-western, Itcha-Ilgachuz, Central-eastern, Jasper-Banff and South-eastern (see Table 1 for details).

**Figure 1:**
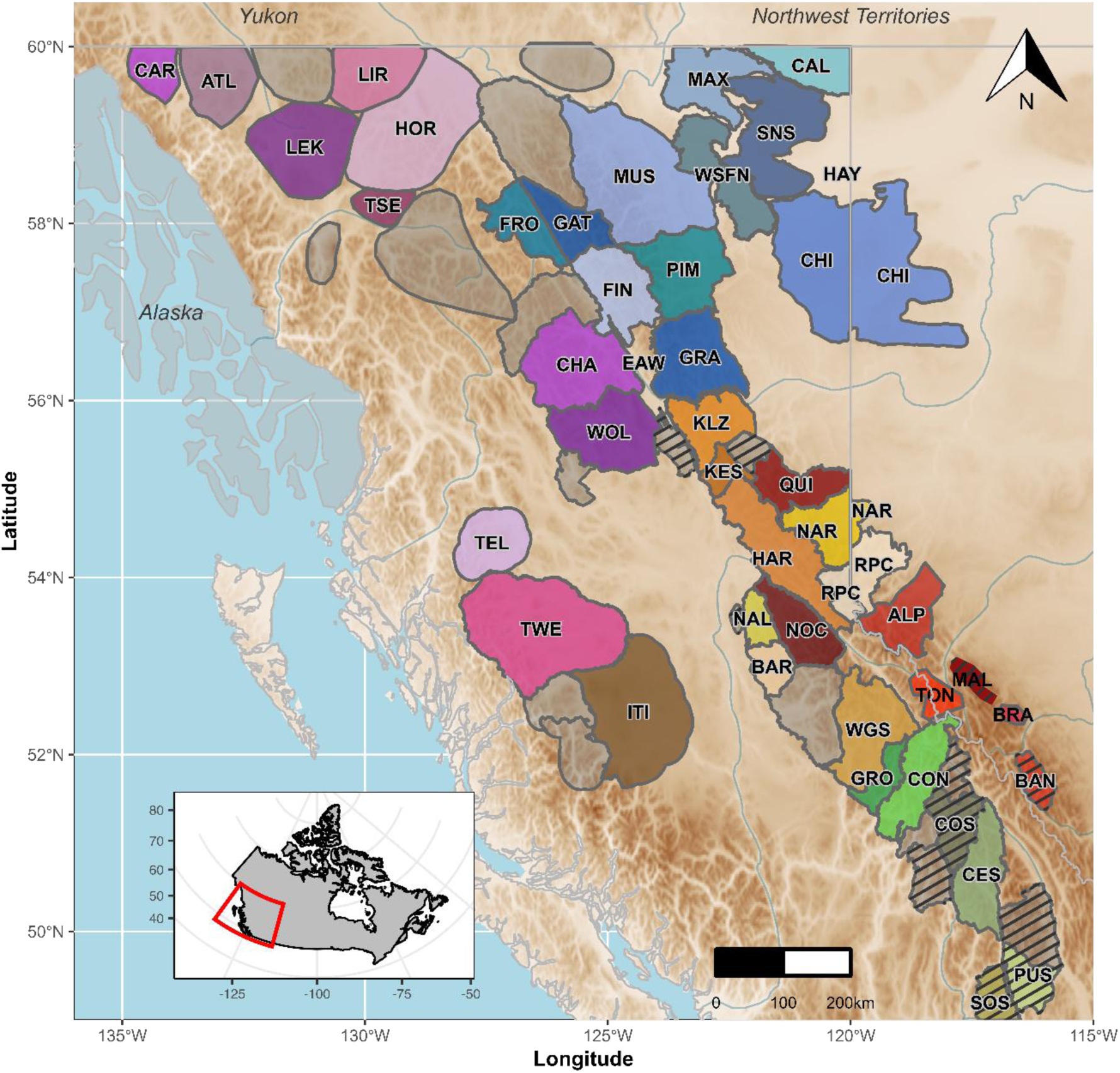
Map of 45 caribou (Rangifer tarandus) subpopulations studied in western Canada. Subpopulations in grey were not studied, and those with striped patterns are considered extirpated, as reported by the Government of British Columbia (2025) and Parks Canada (2023). Full names and details of subpopulations and metapopulations are provided in Table 1. Map created using R packages *ggspatial*, *hydroTSM*, *elevatr*, *ggsn* and *sf*.

**Table 1:**
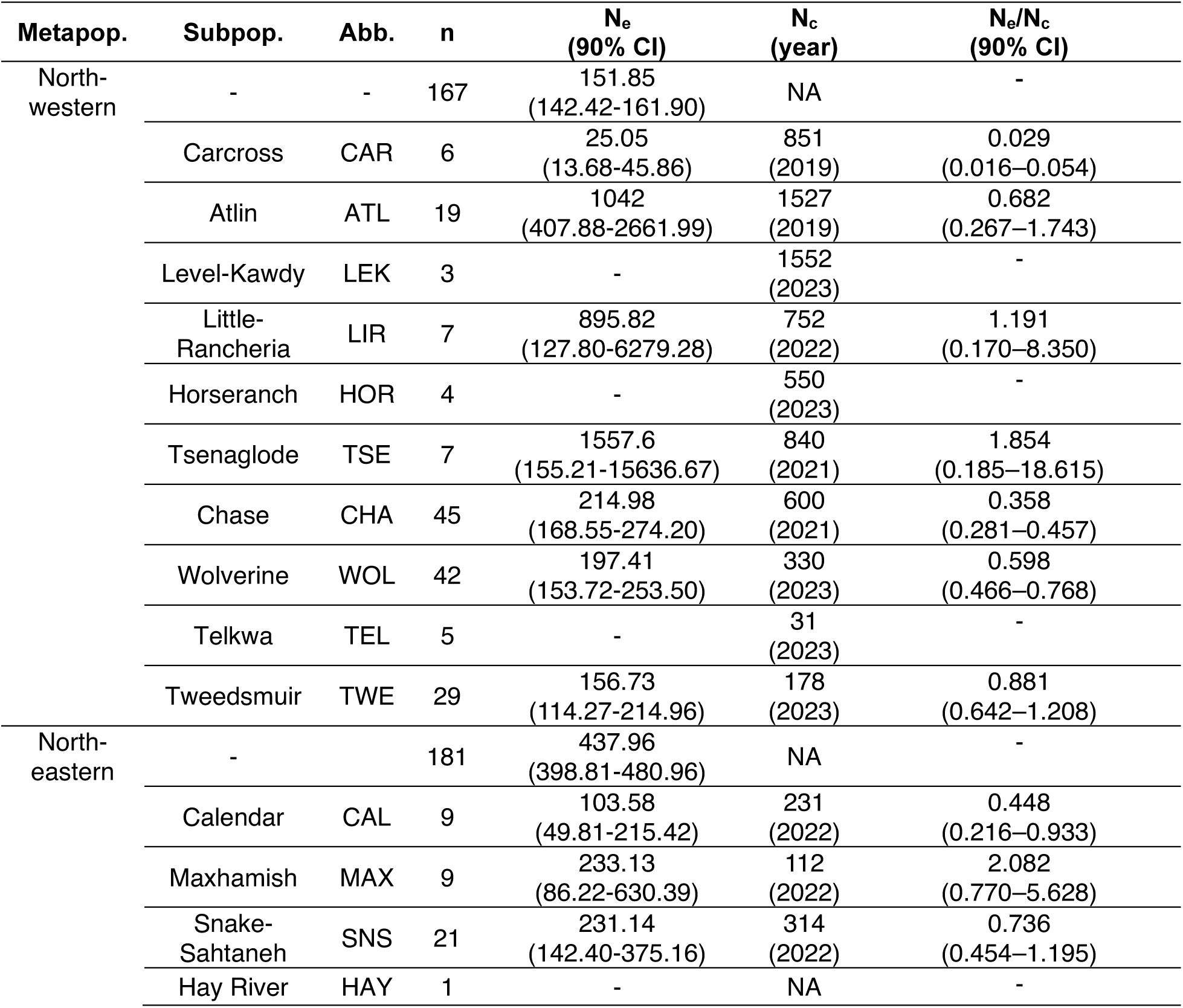
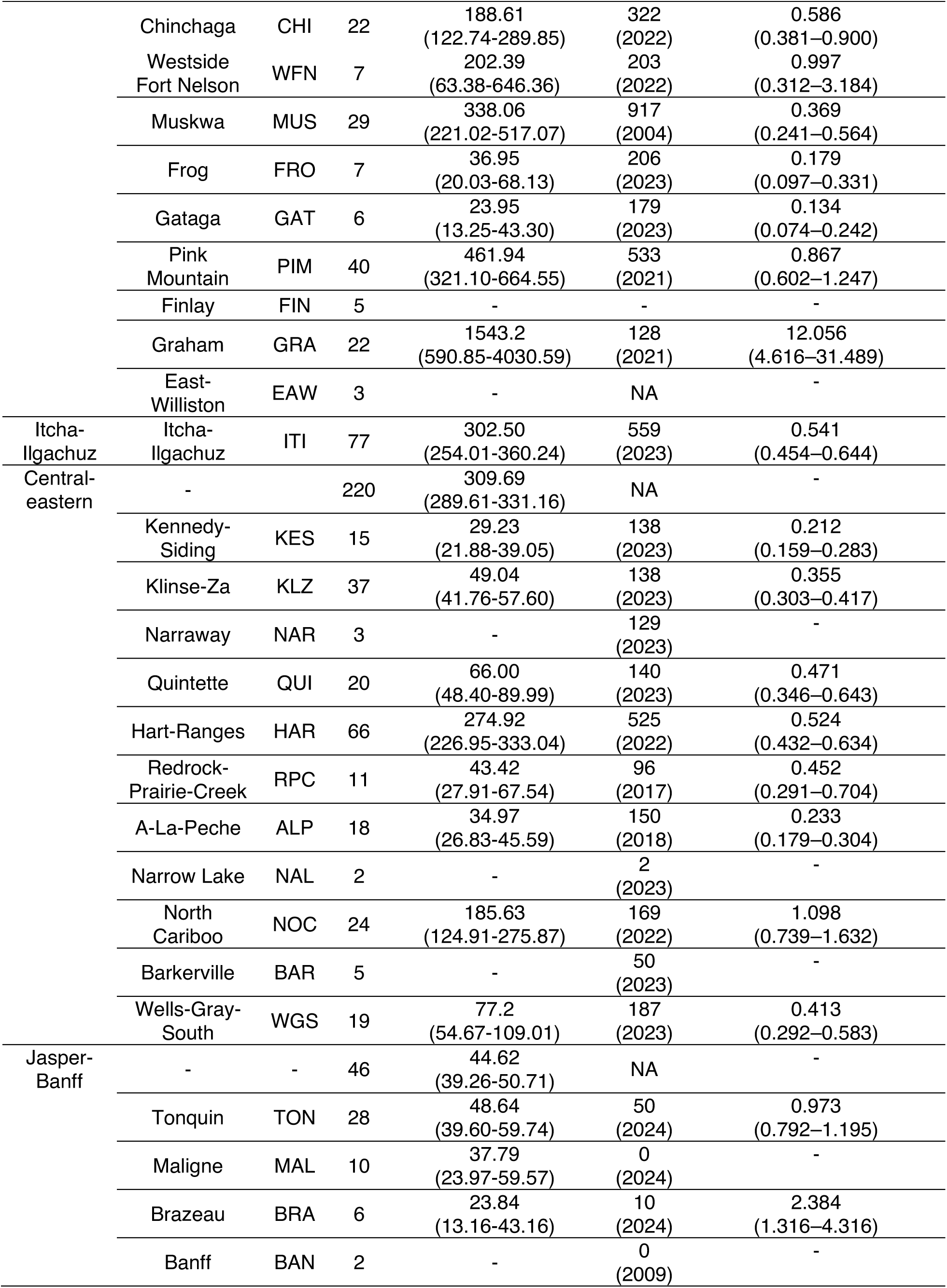
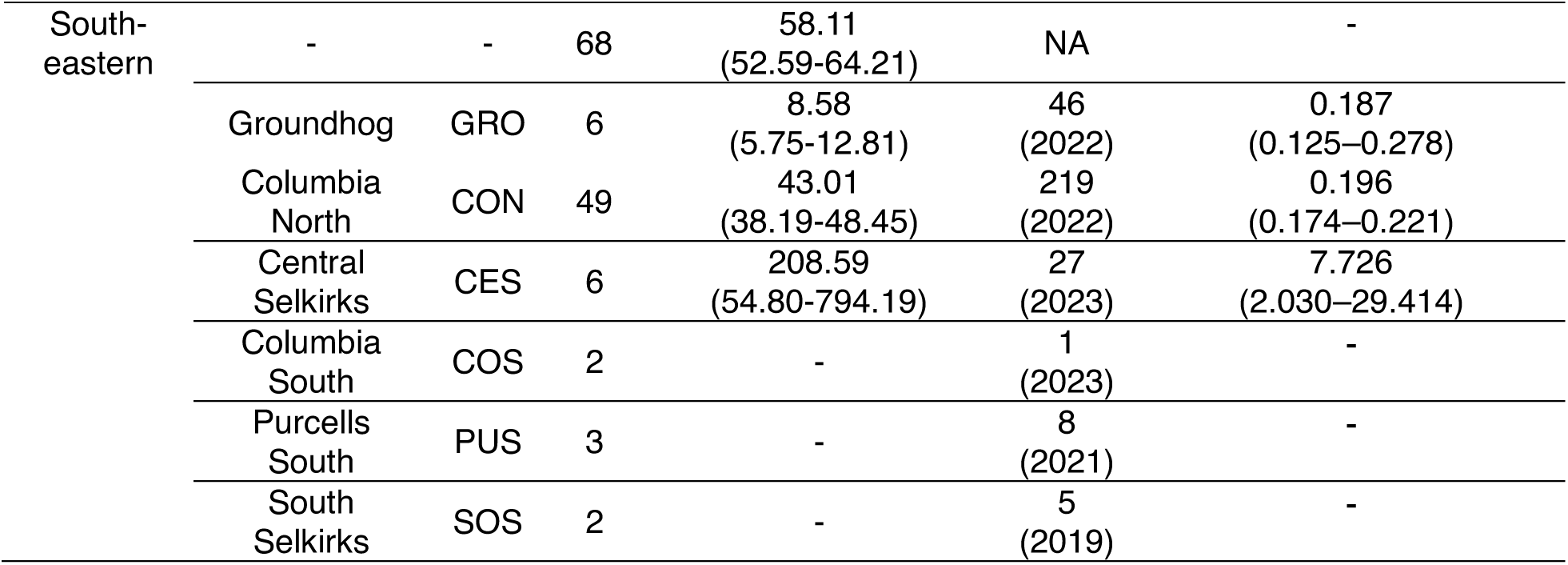
Studied caribou (Rangifer tarandus) subpopulations and metapopulations in western Canada, along with their estimated contemporary effective (N_e_) and census (N_c_) population sizes. Metapopulation clustering followed Deakin et al. (2025). A total of 759 samples from 45 subpopulations were analyzed after filtering (see Materials and Methods). Contemporary demographic estimates include N_e_, calculated using currentNe2 (Santiago et al., 2025), and the N_e_/N_c_ ratio, where N_c_ is based on subpopulation-specific estimates provided by the Government of British Columbia (2025), the Government of Alberta (2017), and Parks Canada (2018, 2023). The year of the most recent N_c_ estimate is indicated in brackets. The 90% confidence interval (CI) for N_e_ was determined using jackknife resampling. Dash symbols (–) indicate values that were not estimable (*e.g.* N_e_ for *n* ≤ 5) while NA indicates values which were not available. Abbreviations: Abb., abbreviation; Metapop., metapopulation; Subpop., subpopulation.

DNA was extracted with QIAGEN DNeasy Blood & Tissue and QIAamp 96 DNA QIAcube HT kits and quantified with a BioTek Synergy LX Multimode Reader (Thermo Fisher Quant-iT dsDNA Assay Kit) or Thermo Fisher Qubit 4 Fluorometer (Qubit dsDNA Assay Kit). Subsequently, 400 ng of DNA per sample was dried and shipped to *Centre d’expertise et de service Génome Québec* (Montreal, Canada) for genotyping using an Illumina SNP array designed for caribou (Carrier et al., 2022). This array provides information for ∼63,000 SNPs evenly distributed along the caribou genome (Carrier et al. 2022).

### Quality control and chromosome assignment

First, we excluded SNPs that were included on the array but found to have poor genotyping rates (3,966) or to be monomorphic (8,749) by Carrier et al. (2022), leading to a total of 63,335 SNPs. Second, as in Deakin et al. 2025, we remapped SNPs from the scaffold-level genome assembly used for array design (ULRtarCaribou_2; GCA_019903745.1) to the newest chromosome-level assembly available (ULRtarCaribou_2v2; GCA_019903745.2) using bowtie2 v2.3.0 (Langmead & Salzberg, 2012). Default parameters were used, except for allowing a maximum of one mismatch during the seed alignment step (-N 1) to allow for slight variation. Next, we used SAMtools v1.18 (Li et al., 2009) to process the alignment data, including filtering, sorting mapped reads, and mapping SNPs to the reference genome. SNPs without a unique chromosome-level position in the newer assembly were excluded. The remapped dataset contained 48,588 SNPs distributed across 35 chromosomes.

After having updated genomic positions, genotypes were filtered using PLINK v1.9 (Purcell et al., 2007). We first excluded 19 samples identified as duplicates by identifying pairs of individuals with an identical by state value greater than 0.95 with the *--genome* option. We then successively removed loci and samples with more than 5% missing data (*--geno* 0.05 and *--mind* 0.05), loci that deviated significantly from Hardy-Weinberg equilibrium (*p <* 1e-05, *--hwe* 0.00001), and duplicate SNPs (*--list-duplicate-vars)*. Following Meyermans et al. (2020), we did not apply filtering based on minor allele frequency or linkage disequilibrium (LD), as such filtering can bias ROH detection and LD-based N_e_ estimates by removing informative rare variants and reducing the linkage signal necessary for accurate inference. This filtering resulted in 33,531 loci. Finally, we excluded SNPs located on sex chromosomes, as done in Hewett et al. (2023) and Perez-Gonzalez et al. (2025), because these have different inheritance patterns, N_e_, and recombination rates compared to autosomes (Ellegren, 2009). The resulting dataset contained 33,346 SNPs for 759 unique individuals across 45 subpopulations (Table 1).

### ROH detection and analysis

ROHs were determined for each individual using the R package detectRUNS v0.9.6 (Biscarini et al., 2018). The consecutive rather than the sliding window approach was used because the latter is known to be less accurate and to overestimate the numbers of ROHs (Smaragdov, 2021, 2023). After evaluating the impacts of parameter choices, we chose to set the minimum number of SNPs (*minSNP)* to 20, and both the maximum number of heterozygous SNPs (*maxOppRun)* and maximum number of missing genotypes in a run (*maxMissRun)* to 2, as this parameter combination struck a balance between stringency and sensitivity (Supporting Information S1, Figure S1). The minimum length of a run was set at 300 kb, and the maximum gap between consecutive SNPs was set to 1 Mb. For each individual, we calculated the number (N_ROH_), average length (L_ROH_), and total length (S_ROH_) of ROHs.

We used Kruskal-Wallis tests to determine whether subpopulations and metapopulations differed in their median N_ROH_ or L_ROH_ values. In addition to reporting the significance of these differences, we calculated effect sizes using epsilon-squared ( ε²) with the formula ε² = (H – k + 1) / (n – k), where H is the Kruskal-Wallis χ² statistic, k is the number of groups, and n is the total sample size. We also used one-sample Wilcoxon signed-rank tests to assess whether each subpopulation ’ s or metapopulation ’ s N_ROH_ and L_ROH_ values differed significantly from the overall median.

### Estimation of inbreeding coefficients

We estimated individual inbreeding coefficients with ROHs (F_ROH_) by dividing S_ROH_ by the length of the genome covered by the SNPs included in the analysis (1,984,467,451 bp), using the *Froh_inbreeding()* function from detectRUNS. We also estimated the contribution of ROHs of different length categories to F_ROH_ for each individual, subpopulation, and metapopulation using the function *Froh_inbreedingClass()*, which we modified to accommodate custom length categories: short (0.3-2 Mb), short-medium (2-4 Mb), medium (4-6 Mb), medium-long (6-8 Mb), and long (> 8 Mb). We used Kruskal-Wallis tests to determine if either subpopulations or metapopulations differed in their median F_ROH_ values and calculated effect sizes using ε². When this test was significant, we used Dunn ’ s Multiple Comparisons test implemented in the R package FSA v0.9.5 (Ogle & Ogle, 2017) to identify pairs that differed. In a separate analysis, we used one-sample Wilcoxon signed-rank tests to assess whether each subpopulation’s F_ROH_ values differed significantly from the overall median, identifying which subpopulations had higher or lower inbreeding levels relative to the entire dataset.

Second, we estimated individual kinship (𝜙_i_) and kinship between pairs of individuals (𝜙_ij_) with the *inbr()* function from the R package popkin v1.3.23 (Ochoa & Storey, 2021). Individual kinship reflects the probability of homozygosity by descent within a single individual, whereas kinship between pairs of individuals reflects the probability that an allele in one individual is identical by descent (IBD) to an allele in another individual. The *inbr()* function also provides inbreeding coefficients calculated from individual kinship estimates (F_kin_ = 2𝜙_i_ - 1, where F_kin_ represents the inbreeding coefficient; Ochoa & Storey, 2021). We examined the relationship between F_kin_ and F_ROH_ using two complementary approaches. First, we performed Standardized Major Axis (SMA) regression, which accounts for measurement error in both variables, to test whether the relationship approximated the theoretical 1:1 expected when both metrics quantify the same biological phenomenon. For subpopulation-level comparisons, we used ordinary least squares (OLS) regression, as SMA slope estimates become unreliable with the small sample sizes (*n* < 5) present in some subpopulations. Second, we used Bland-Altman analysis to assess the agreement between F_kin_ and F_ROH_ by plotting the difference between the two measures (F_kin_-F_ROH_) against their mean and calculated the mean difference (bias) and standard deviation of differences for each subpopulation.

### Contemporary N_e_ and N_e_/ N_c_ ratio

We estimated contemporary N_e_ using currentNe2 v2.0 (Santiago et al., 2025), applying a genome-wide LD integration approach that accounts for physical distances between markers and population-specific mating systems. Since caribou-specific recombination rates are unknown, we used *r* = 1.0 cM/Mb for main results (the typical mammalian rate; Dumont & Payseur, 2008) and tested sensitivity at *r* = 0.9 and 1.1 cM/Mb (Crossman et al., 2024; results in Tables S3-S4). The expected number of full siblings per individual (*k*) was estimated from the data using default parameters.

currentNe2 includes a structure correction option (-x) for metapopulations, assuming an island model with equal-sized subpopulations and symmetrical migration. Caribou violate these assumptions: subpopulations vary 100-fold in size (Table 1), exhibit hierarchical rather than island structure (Deakin et al., 2025), and show asymmetric migration (Theoret et al., 2022). While Santiago et al. (2025) demonstrate that the -x option provides robust N_e_ estimates even under substantial assumption violations, the method is optimized for scenarios with moderate differentiation (N_m_ ≈ 1-10) and can produce less stable parameter estimates when applied to systems with extreme size asymmetries and complex hierarchical structure. We therefore used the panmictic model, which provides stable subpopulation-specific N_e_ estimates with well-defined confidence intervals (CIs, determined using jackknife resampling) when sampling is restricted to single demographic units (Santiago et al., 2025). N_e_ was not estimated for 23 subpopulations with small sample sizes (*n* ≤ 5).

We calculated N_e_/N_c_ ratios by dividing the contemporary N_e_ point estimate by the most recent N_c_, with 90% confidence intervals derived by dividing the lower and upper bounds of the N_e_ jackknife 90% CIs by the N_c_ point estimate, treating N_c_ as fixed. N_c_ estimates were obtained from the Government of British Columbia (2025), the Government of Alberta (2017), and Parks Canada (2018, 2023) and derive from Integrated Population Modelling (IPM), observed total or sample-based counts, minimum number known alive, and expert knowledge. We compared N_e_/N_c_ ratios to established thresholds (0.1 for potential genetic vulnerability; Frankham 1995). For all demographic analyses, we included only the 31 subpopulations with sample sizes *n* > 5 and available census estimates (N_c_ ≠ 0).

### Relation between inbreeding and population size metrics

Using the subset of 31 subpopulations used in the N_e_/N_c_ ratios analysis, we examined the relationship between mean F_ROH_ and population metrics (N_c_ and N_e_) using linear models with log-transformed F_ROH_ as the response variable. Candidate models included log-transformed N_c_, N_e_, their quadratic terms, and interactions to test for linear, non-linear, and combined effects of population size metrics.

Model selection was performed using second-order Akaike Information Criterion (AICc) implemented in the R package MuMIn v1.48.4 (Bartoń, 2016). Models with ΔAICc < 2 were considered to have substantial support and were used for model averaging. We performed conditional model averaging (revised.var = TRUE, full = FALSE) using the model.avg() function from MuMIn, which averages parameter estimates only across models where each parameter appears, providing more reliable estimates for prediction (Grueber et al., 2011). For the top-ranked models, we calculated the percent change in F_ROH_ associated with a 10-fold increase in either N_c_ or N_e_ using the formula (10*^β^*-1) × 100%, where *β* is the estimated regression coefficient. Model predictions were back-transformed to the original scale and plotted with 95% CIs alongside observed data. When interaction terms were present in supported models (ΔAICc < 2), we visualized the interaction effects by calculating predicted F_ROH_ across a range of N_c_ values at fixed N_e_ levels (5, 10, 25, 50, 100, 250, 500). All analyses were conducted in R version 4.2.3 (R Core Team, 2024)

### Historical N_e_ and demographic history

We reconstructed historical N_e_ using GONE2 v2.0 (Santiago et al., 2025), which estimates temporal N_e_ changes from LD patterns across recombination bins. As for contemporary N_e_, we used *r* = 1.0 cM/Mb (main results) with sensitivity analyses at *r* = 0.9-1.1 cM/Mb, and the panmictic model. To obtain robust estimates, we ran each analysis with 100 independent seeds and report mean N_e_ ± SD across runs. Chromosome 33 was excluded from these analyses as its genetic length fell below the 20 cM threshold required for reliable N_e_ estimation, resulting in 31,859 SNPs for this analysis. Generation time was assumed to be 8 years (mean of 7-9, Government of British Columbia, 2025) when converting generations to calendar years. As for currentNe2, GONE2 analyses were limited to 32 subpopulations with *n* > 5.

In addition to GONE2 analyses, we compared four demographic scenarios using Approximate Bayesian Computation with Random Forests (DIYABC-RF v1.0; Collin et al., 2021), testing four models: (1) constant N_e_, (2) recent decline (within 5 generations), (3) old decline (25-100 generations ago), and (4) both old and recent declines. We assigned uniform prior distributions to all demographic parameters, with ranges selected to encompass a broad spectrum of possible demographic histories: *N0* = 10-1,000, *N1/N2* = 10-10,000, *t1* = 1-5 generations (∼8-40 years), and *t2* = 25-100 generations (∼200-800 years). Scenario support was evaluated using chi-square tests, pairwise proportion tests with FDR correction, and Cramer’s *V*. Details on data preparation and parameter justification are provided in Supporting Information S2.

## Results

### Numbers and lengths of ROHs

Mean ROH number (N_ROH_, Figure S3) was 71.19 ± 25.44 (median = 63) and mean ROH length (L_ROH_, Figure S4) was 2.55 ± 0.89 Mb (median = 2.36). Subpopulations differed significantly in both metrics (Kruskal-Wallis, both *p <* 0.001), with subpopulation identity explaining 72.9% of N_ROH_ variation (*ε*² = 0.73) and 39.8% of L_ROH_ variation (*ε*² = 0.40).

Thirteen subpopulations showed significantly lower N_ROH_ than the overall median (one-sample Wilcoxon signed-rank test, all *p <* 0.05, Figure S3), while Tweedsmuir, Itcha-Ilgachuz, Wells Gray South, Columbia North, and the Jasper-Banff group were significantly higher. For L_ROH_, 16 subpopulations showed significantly lower values than the overall median (one-sample Wilcoxon signed-rank test, all *p* < 0.05, Fig. S4). Only Wells Gray South and Columbia North were significantly higher than the median.

### Variation in inbreeding coefficients (F_ROH_)

Mean F_ROH_ was 0.096 ± 0.059 (median = 0.075, range: 0.016-0.346), and differed significantly among subpopulations (Kruskal-Wallis, *p <* 0.001, *ε*² = 0.43). Inbreeding showed a strong south-to-north gradient (Figure 2A). The highest F_ROH_ values occurred in southern and central subpopulations: Purcells South (0.302 ± 0.023), Banff (0.284 ± 0.001), Columbia South (0.227 ± 0.123), and Itcha-Ilgachuz (0.203 ± 0.030). Northern subpopulations generally exhibited low inbreeding, including Frog (0.039 ± 0.012), Tsenaglode (0.045 ± 0.012), and Little Rancheria (0.046 ± 0.012). Relative to the range-wide median, 11 subpopulations (predominantly southern) showed significantly elevated F_ROH_, while 16 subpopulations (predominantly northern) showed significantly reduced F_ROH_ (Wilcoxon tests, all *p <* 0.05; Figure 2B, Table S1). Mean F_ROH_ exceeded 0.1 in 14 subpopulations.

**Figure 2:**
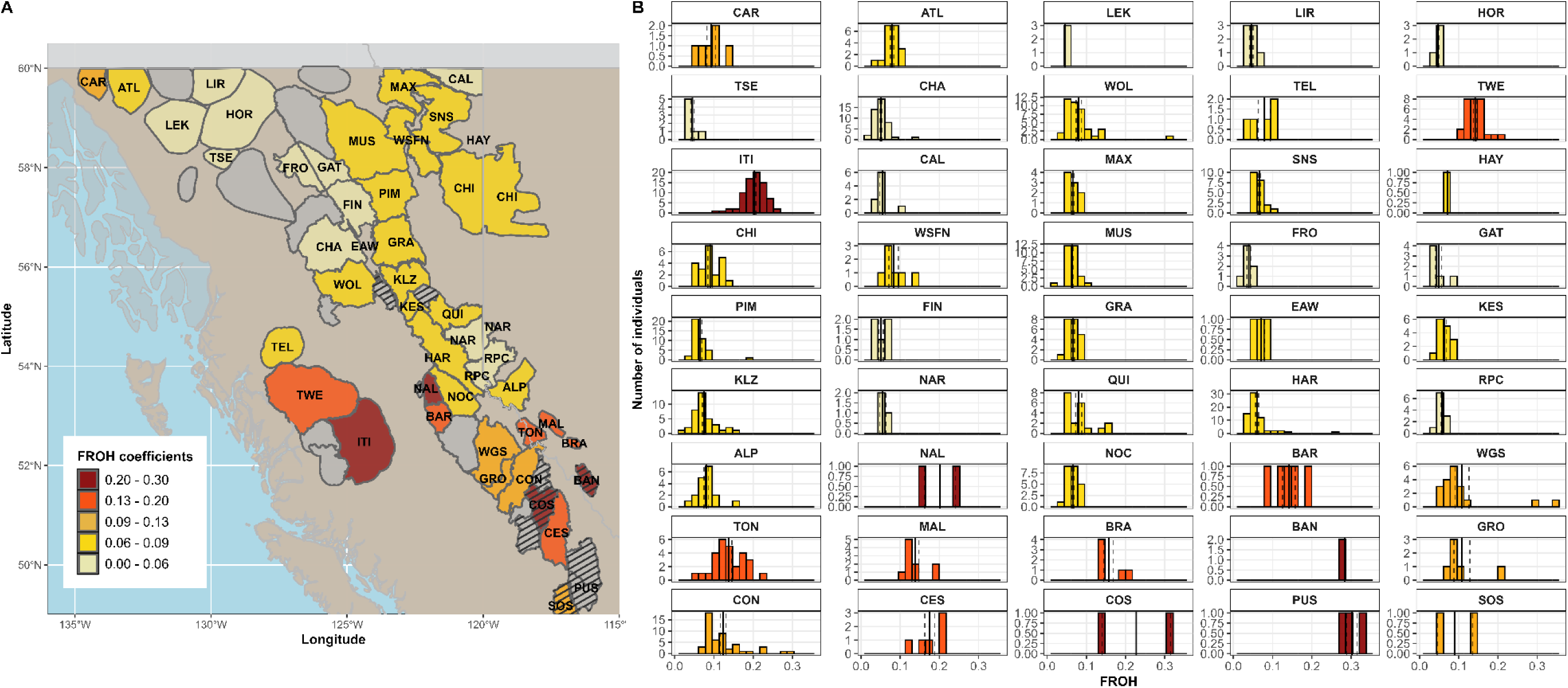
Distribution of inbreeding coefficients derived from Runs of Homozygosity (F_ROH_) for 45 caribou (Rangifer tarandus) subpopulations in western Canada. A. Map showing the spatial distribution of sampled subpopulations with colour-coding based on inbreeding coefficient (F_ROH_) values. Subpopulations are labelled with abbreviations detailed in Table 1. Map created using R packages *ggsn*, *ggspatial* and *sf*. B. Histograms showing the distribution of individual F_ROH_ values within each subpopulation. Colours of the bars are related to their colours on the map. Solid vertical lines correspond to the mean and dotted lines to the standard deviation.

F_ROH_ varied significantly among metapopulations (Kruskal-Wallis, *p <* 0.001, *ε*² = 0.43, Figure S5). The Itcha-Ilgachuz, Jasper-Banff (0.147 ± 0.047), and South-eastern metapopulations (0.137 ± 0.066) showed the highest mean F_ROH_, while the North-eastern metapopulation exhibited the lowest (0.067 ± 0.022). Dunn’s Tests identified pairs of metapopulations significantly differing from each other, even after Bonferroni correction, notably the Itcha-Ilgachuz and Central-eastern metapopulations, with the former exhibiting substantially higher inbreeding (Dunn’s Test, all *p <* 0.001). Other notable differences included higher F_ROH_ in the South-eastern compared to the North-western metapopulation (*Z* = -7.53, *p <* 0.001), and lower F_ROH_ in the North-eastern compared to the Jasper-Banff metapopulation (*Z* = 8.70, *p <* 0.001).

### ROH length classes

ROH length classes contributed differently to inbreeding (Kruskal-Wallis, *p <* 0.001, *ε*² = 0.34; Figure 3), with all pairwise comparisons being significant (Dunn’s tests, all *p <* 0.001). Long ROHs (> 8 Mb) contributed most to overall F_ROH_ (0.035 ± 0.038), followed by short ROHs (< 2 Mb; 0.033 ± 0.010). Medium-length ROHs showed progressively smaller contributions: 2-4 Mb (0.019 ± 0.012), 4-6 Mb (0.012 ± 0.010), and 6-8 Mb (0.009 ± 0.009).

**Figure 3:**
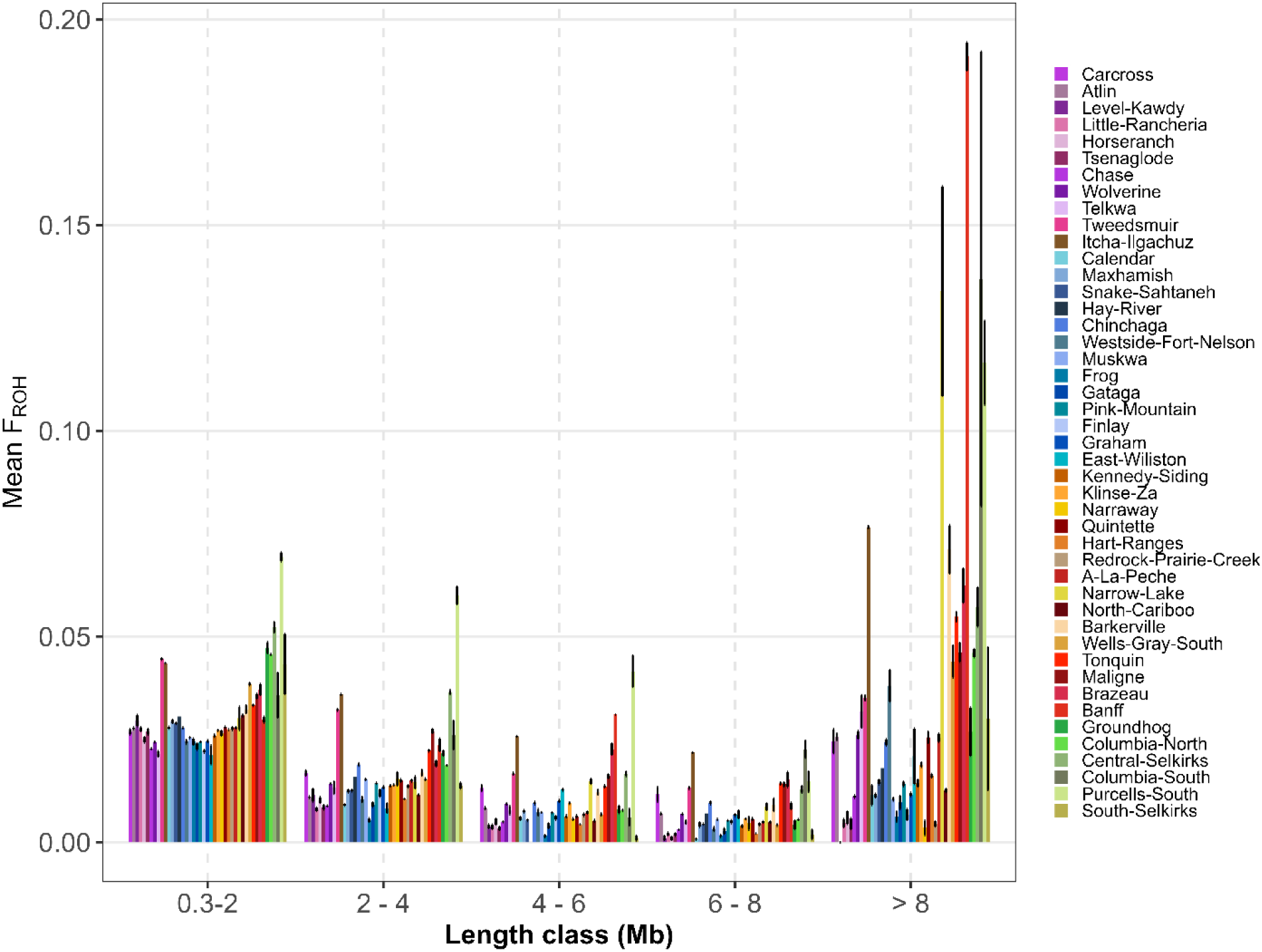
Inbreeding coefficient derived from Runs of Homozygosity (F_ROH_) per class of ROHs lengths for 45 subpopulations of caribou (Rangifer tarandus) in western Canada. The different classes are short: 0.3 – 2 Mb, short-medium: 2-4, medium: 4-6, medium-long: 6-8 Mb, long: > 8 Mb. The standard error is represented for each subpopulation.

The relative contribution of different ROH length varied across regions (Figure 3). Southern subpopulations (Maligne, Banff, Columbia South, Purcells South) showed higher contributions from long ROHs (> 8 Mb) relative to other length classes, while northern subpopulations exhibited more balanced contributions across length categories. At the metapopulation level, Itcha-Ilgachuz, Jasper-Banff and South-eastern metapopulations had the highest F_ROH_ values for long ROHs (0.076, 0.054 and 0.046, respectively; Figure S6). The North-eastern metapopulation exhibited the lowest F_ROH_ values across most length categories.

### Comparison of inbreeding metrics

F_ROH_ showed strong concordance with kinship-based inbreeding estimates (F_kin_; *r²* = 0.96, *p <* 0.001, Figure S7). The SMA slope was 1.035 (95% CI: 1.020-1.049), slightly but significantly greater than the theoretical 1:1 relationship, indicating that F_kin_ values were marginally higher than F_ROH_ values, particularly at higher inbreeding levels. Bland-Altman analysis confirmed this pattern, with F_kin_ generally exceeding F_ROH_ across all subpopulations (mean difference = 0.014 ± 0.012; Figure S8). At the subpopulation level, most groups showed slopes near or above one using OLS regression though some exhibited markedly different patterns, particularly those with small sample sizes (e.g., Level-Kawdy: slope = -2.57, *n* = 3; Horseranch: slope = -0.23, *n* = 4).

### Contemporary N_e_ and N_e_/ N_c_ ratio

Contemporary N_e_ was estimated for 32 subpopulations (Table 1) and ranged from 8.58 (Groundhog) to 1557.60 (Tsenaglode). Most subpopulations (28) had N_e_ point estimates < 500 (the long-term viability threshold), with 22 having 90% CIs entirely below 500. Twelve subpopulations had N_e_ point estimates < 50, with seven having CIs entirely below 50 (A La Peche, Carcross, Gataga, Kennedy-Siding, Brazeau, Groundhog, and Columbia North), indicating short-term vulnerability. Graham was the only subpopulation with both an N_e_ point estimate and a lower CI bound exceeding 500.

N_e_ estimates were sensitive to the assumed recombination rate, with values at *r* = 0.9 and *r* = 1.1 cM/Mb averaging 3% higher and 3% lower, respectively, compared to *r* = 1 (Table S2). Despite these shifts, the relative ranking among subpopulations and the pattern of subpopulations above or below critical thresholds remained consistent across recombination rates.

At the metapopulation level, N_e_ point estimates ranged from 44.62 (Jasper-Banff) to 437.96 (North-eastern), with generally narrower CIs indicating higher precision at this broader scale (Table 1). All six metapopulations had N_e_ point estimate and higher 90% confidence bounds < 500. Jasper-Banff was the only metapopulation with N_e_ point estimate < 50; although its 90% CI included this threshold (Table 1). The North-eastern, Central-eastern, and Itcha-Ilgachuz metapopulations exhibited the highest values (N_e_ > 300). Similar to subpopulations, metapopulation estimates showed consistent sensitivity to assumed recombination rates (Table S3).

N_e_/N_c_ ratios were estimable for 31 subpopulations and showed substantial variation, ranging from 0.029 (Carcross) to 12.056 (Graham) with an overall median of 0.462 (Table 1). The majority of these ratios (23 of 31) fell between 0.1 and 1.0. One subpopulation showed a ratio below 0.1 (Carcross: 0.029), while seven subpopulations exceeded 1.0 (Little-Rancheria, Tsenaglode, Maxhamish, Graham, North Cariboo, Brazeau, and Central Selkirks). The seven subpopulations with ratios > 1.0 generally exhibited lower sample sizes and wider CIs (with the exception of Graham and North Cariboo) compared to those with ratios between 0.1 and 1.0.

### Relationships between F_ROH_, N_e,_ and N_c_

N_c_ was the primary predictor of F_ROH_, whereas contemporary N_e_ showed minimal predictive power (Table S4). Model selection identified the linear N_c_ model as best supported (*ΔAICc* = 0, *w* = 0.43), followed by the quadratic N_c_ model (*ΔAICc* = 0.21, *w* = 0.38), with model-averaged predictors illustrating a consistent decrease in F_ROH_ with increasing N_c_ (Figure 4A). In the top-ranked model, the effect of N_c_ was significant (*β* = -0.173, *p* = 0.015, *R^2^* = 0.19; Table S5). The quadratic model explained more variance (*R^2^*= 0.252), though its individual linear (*β* = -0.78, *p* = 0.069) and quadratic (*β* = 0.059, *p* = 0.148) terms were not strictly significant. Accordingly, the model-averaged estimate for N_c_² was also not statistically significant (Table S5). In contrast, models including N_e_ received minimal support (*ΔAICc* > 2.6, combined weight < 0.19), and the N_e_-only model explained only 5% of the variance (*R²* = 0.05, *ΔAICc* > 4.98, *w* < 0.04, Figure 4B).

**Figure 4:**
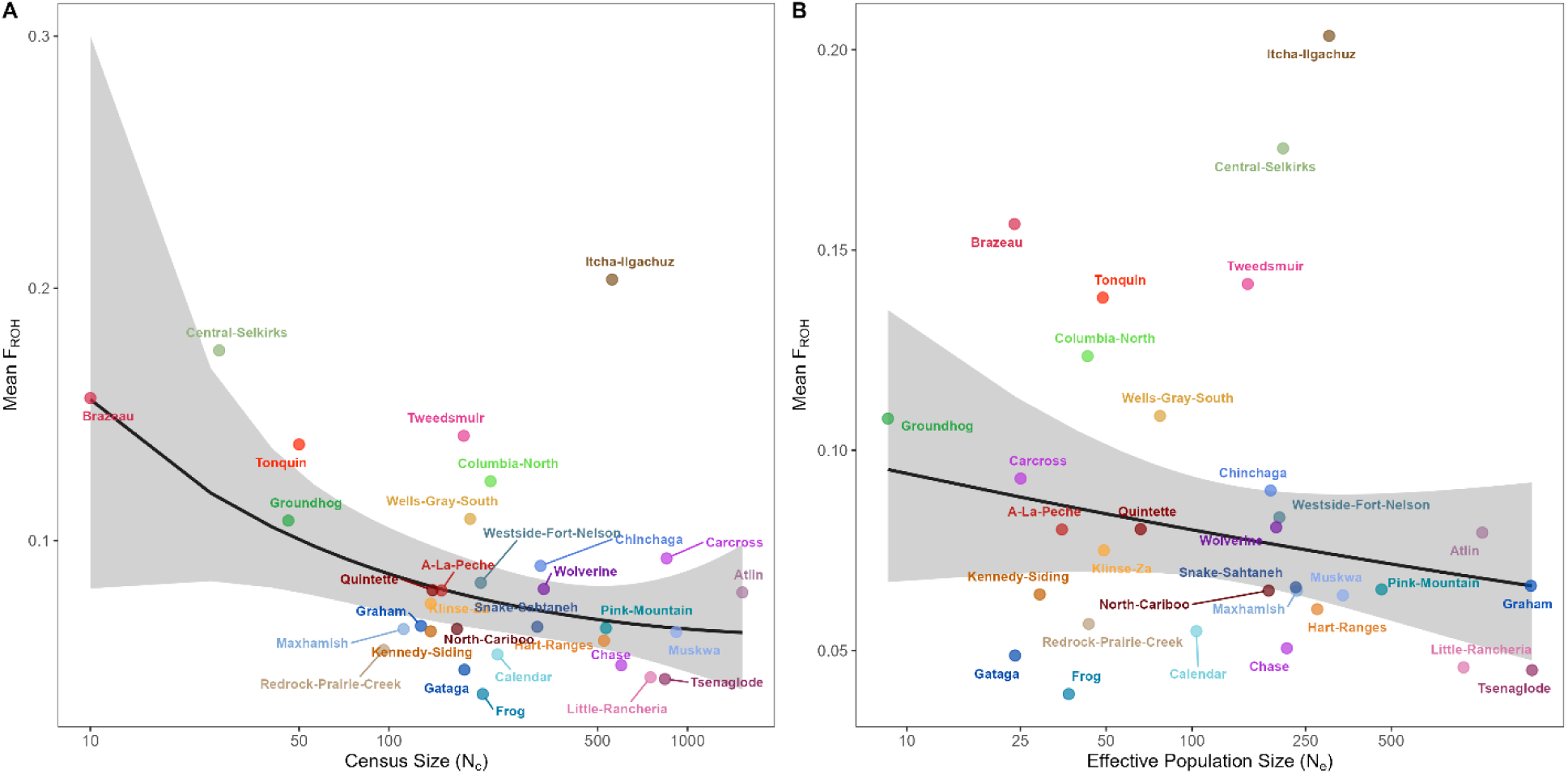
Relationship between census population size (N_c_), contemporary effective population size (N_e_) and inbreeding coefficient derived from Runs of Homozygosity (F_ROH_) for 31 caribou (Rangifer tarandus) subpopulations in western Canada. Panels show the relationship between (A) N_c_ and mean F_ROH_; (B) N_e_ and mean F_ROH_. Each point represents a subpopulation. Solid lines show model predictions with 95% confidence intervals (shaded areas). Panel A shows model-averaged predictions from the top two models; Panel B shows predictions from the N_e_-only model. All models were fitted to log-transformed data and back-transformed for visualization. Subpopulations with small sample sizes (*n* ≤ 5) or N_c_ = 0 (Maligne) were excluded from analyses.

### Historical N_e_

Historical N_e_ reconstruction revealed a north-to-south temporal gradient in bottleneck timing (Figure 5). Northern subpopulations and Itcha-Ilgachuz experienced declines earliest (40-30 generations ago, ∼1700-1780), followed by Central-eastern subpopulations (30-20 generations ago, ∼1780-1860), and most recently southern subpopulations (20-10 generations ago, ∼1860-1940). Most subpopulations showed demographic trajectories concordant with their respective metapopulations, indicating shared demographic histories within regions.

**Figure 5:**
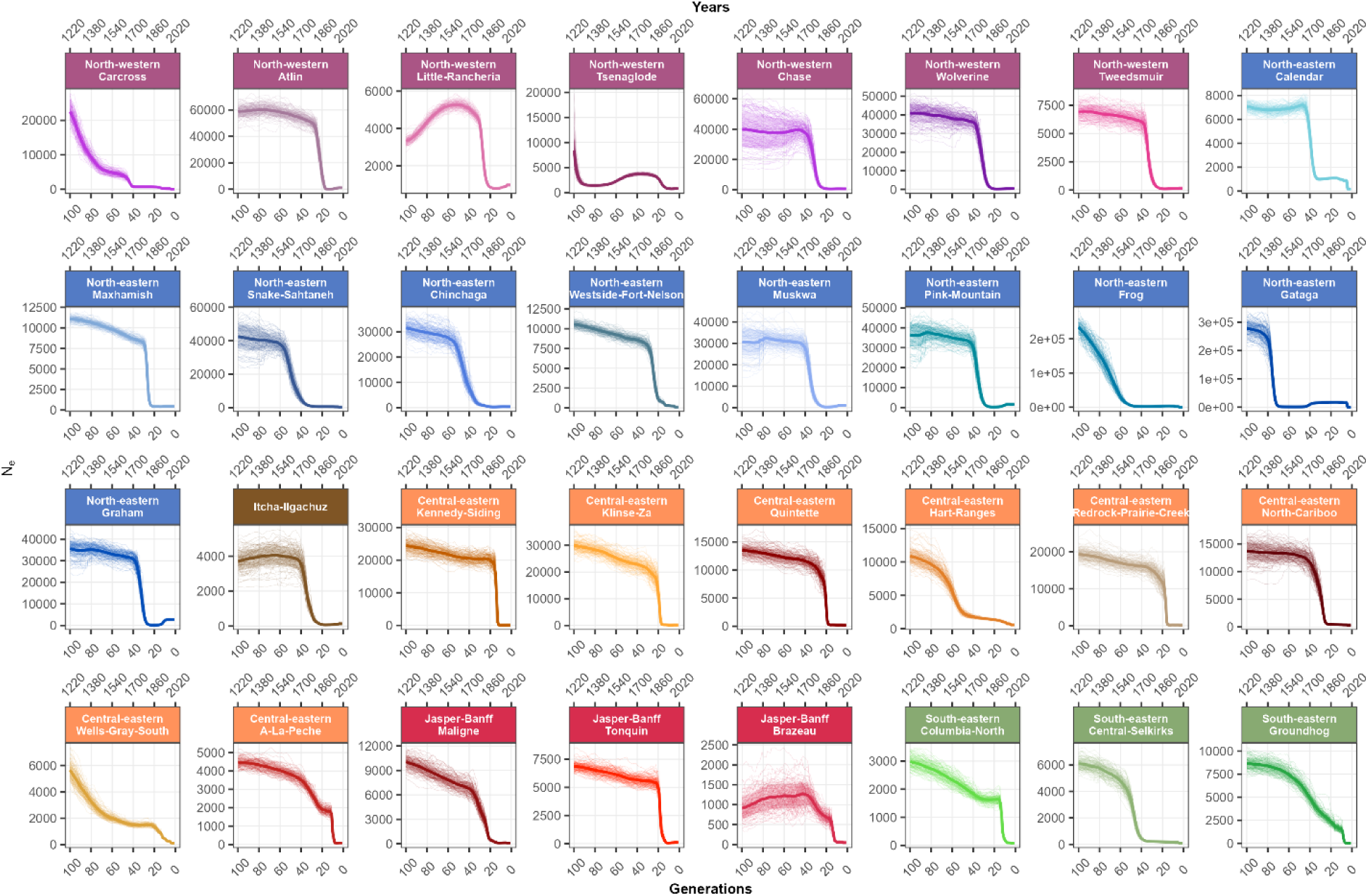
Historical effective population size (N_e_) trajectories over the last 100 generations (∼800 years) for 32 caribou (Rangifer tarandus) subpopulations in western Canada. N_e_ point estimates were obtained using GONE2 v2.0 under a panmictic model. The thick solid lines represent the mean N_e_ estimates calculated from 100 independent stochastic runs (seeds) at a recombination rate of 1.0 cM/Mb. Shaded ribbons represent ± 1 standard deviation around the mean. Subpopulations are grouped and colored by their corresponding metapopulation as defined by Deakin et al. (2025) based on genetic structure analyses. Subpopulations with small sample sizes (*n* ≤ 5) were excluded from analyses. Generations were converted to years (upper x-axis), assuming a mean generation time of 8 years.

Analysis of the last 10 generations (Figure S10) showed that 13 subpopulations, primarily in the north, experienced modest N_e_ increases 10-5 generations ago (∼1940-1980). However, most southern subpopulations exhibited a sharp decrease approximately 5-4 generations ago (∼1980-1998). Over the final four generations (since ∼1988), N_e_ stabilized at historically low levels across most groups, which may also reflect reconstruction limitations in the most contemporary time steps.

The metapopulation analysis revealed trends similar to those observed in the subpopulation analyses, but with comparatively lower historical N_e_ values. (Figure 6). However, the North-eastern metapopulation showed an older bottleneck (60-70 generations ago), which was only detected in Gataga and Frog in subpopulation analyses. Consistent with the subpopulation analyses, the results suggested recent bottlenecks in Jasper-Banff (20-10 generations ago) and recent N_e_ decreases in all metapopulations (15-5 generations), particularly for South-eastern. N_e_ trends were robust to varying recombination rates (0.9 and 1.1) at the metapopulation level (Figure 6), although subpopulations with smaller sample sizes exhibited greater posterior variance in older generations (Figure S11).

**Figure 6:**
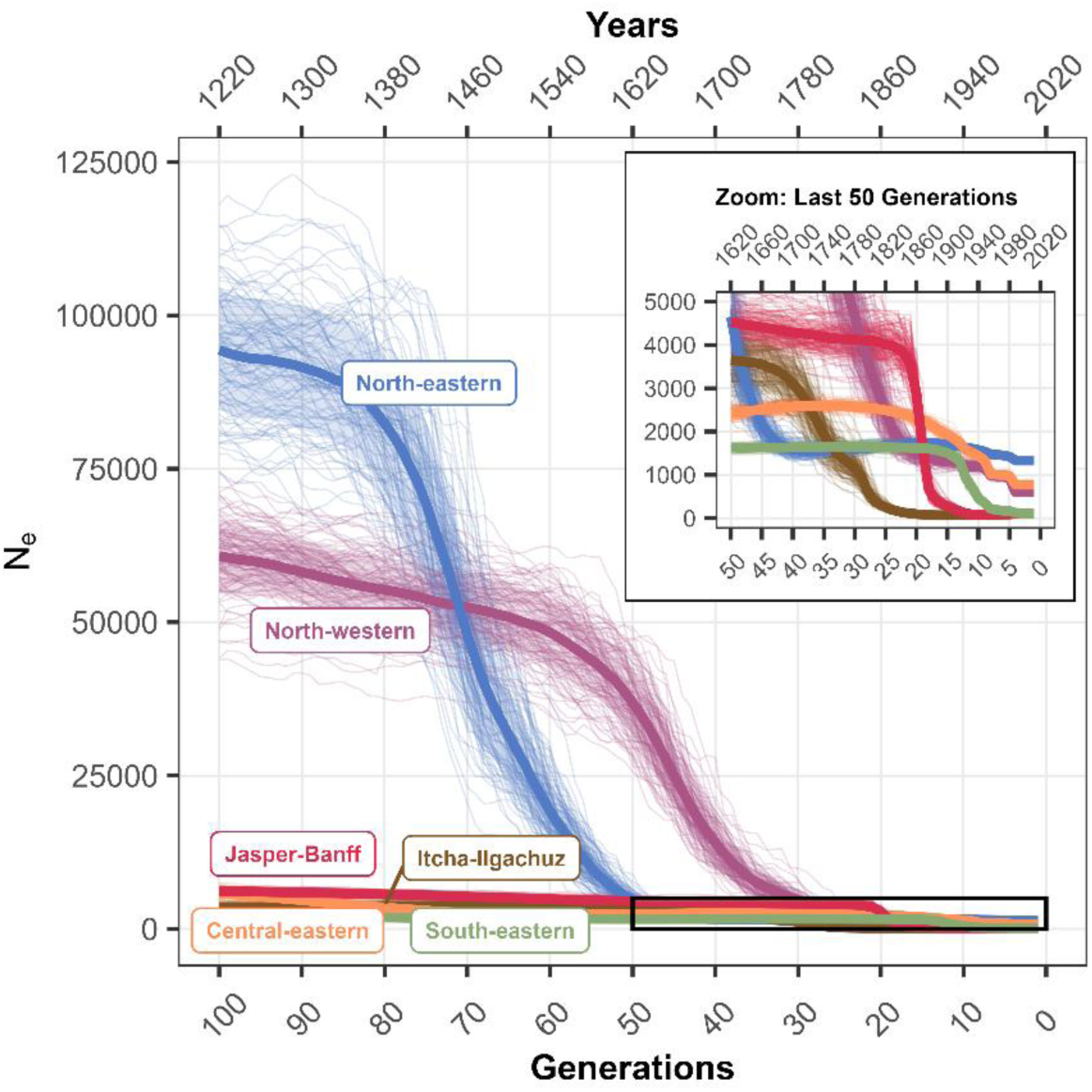
Historical effective population size (N_e_) trajectories over the last 100 generations (∼800 years) for six caribou (Rangifer tarandus) metapopulations in western Canada. N_e_ point estimates were obtained using GONE2 v2.0 under a panmictic model. The thick solid lines represent the mean N_e_ estimates calculated from 100 independent stochastic runs (seeds) at a recombination rate of 1.0 cM/Mb. Shaded ribbons represent ± 1 standard deviation around the mean, while faint background lines display the trajectories of the 100 individual seed runs. The inset shows a magnified view of the most recent 50 generations (∼400 years), corresponding to the period bounded by the black rectangle in the main plot. Metapopulations were defined by Deakin et al. (2025) based on genetic structure analyses. All the samples (*n* = 759) were used for this analysis. Generations were converted to years (upper x-axis), assuming a mean generation time of 8 years.

### Demographic scenarios

Most caribou subpopulations (23 of 32, 71.9%) showed support for an old and recent decline scenario (Figure S12), with posterior probabilities ranging from 0.243 to 0.835 (mean = 0.523, SD = 0.141). For seven subpopulations (21.9%) the old decline scenario was favored, while the recent decline was only supported twice (6.2%). The constant N_e_ scenario received negligible support, with posterior probabilities never exceeding 0.1. Overall, scenario selection deviated significantly from a uniform distribution (*χ*² = 22.562, *p* < 0.001, Cramer’s *V* = 0.594). The old and recent decline scenario was strongly preferred, and all pairwise scenario comparisons were significant (*p* ≤ 0.05).

## Discussion

We assessed inbreeding, N_e_, and demographic history across caribou subpopulations and metapopulations in western Canada using genome-wide SNP data. High inbreeding levels (F_ROH_ > 0.1) were detected in the Jasper-Banff, South-eastern and Itcha-Ilgachuz metapopulations. Multiple subpopulations showed contemporary N_e_ below critical thresholds (N_e_ < 50 and 500).

Historical analyses revealed a north-to-south temporal gradient in bottleneck timing: northern subpopulations declined the earliest (∼1700-1780), followed by central (∼1780-1860) and southern subpopulations (∼1860-1940). These genomic patterns provide insights beyond demographic monitoring by establishing historical baselines that distinguish between earlier environmental shifts and more recent anthropogenic impacts. Ultimately, these results allow us to identify highly vulnerable subpopulations where high inbreeding and low N_e_ signal a need for genetic intervention, even in cases where census sizes might otherwise suggest demographic stability.

Our results indicate that the medium-density SNP array developed by Carrier et al. (2022) provides sufficient resolution for ROH detection across caribou subpopulations. Similar arrays have been used to identify ROH patterns in various species (passerine, *Notiomystis cincta*, Duntsch et al., 2021, 2023; oyster; *Crassostrea gigas*, Gutierrez et al., 2020; cattle; *Bos taurus*, Mastrangelo et al., 2016), particularly for segments exceeding 5 Mb (Ceballos et al., 2018; Szmato ła et al., 2020). Our SNP density limited the detection of very short ROH, which can slightly underestimate F_ROH_ (Bertolini et al., 2018; Dadousis et al., 2021). Due to the parameter-dependent nature of F_ROH_, cross-study comparisons require a consistent methodology to ensure validity. Our analysis revealed a strong positive association between F_kin_ and F_ROH,_ demonstrating that F_ROH_ reliably reflects inbreeding in this system, consistent with findings in other animal species (Marcuzzi et al., 2024; Sweetalana et al., 2025).

Our analysis revealed substantial variation in inbreeding levels. High inbreeding was concentrated in southern subpopulations, with F_ROH_ values exceeding the 0.1 fitness-effect threshold (Rehder et al., 2013) primarily in southern subpopulations, including those in the South-eastern and Jasper-Banff metapopulations that have experienced or are threatened with extirpation (Environment Canada, 2014; Government of British Columbia, 2025), as well as in Itcha-Ilgachuz. These findings provide geographic resolution to the highly variable inbreeding levels previously observed across the CSM lineage (encompassing Central-Mountain, Southern Mountain and Itcha-Ilgachuz herds; Taylor et al., 2024), and complement broader assessments of genetic diversity and inbreeding across these subpopulations (Cavedon et al., 2023; Deakin et al., 2025). Specifically, we show that high inbreeding is concentrated in Southern Mountain and Itcha-Ilgachuz subpopulations, compared to lower levels in Central Mountain and northern subpopulations.

Elevated inbreeding poses a risk of inbreeding depression, for which evidence in caribou remains mixed. While Ralls et al. (1988) documented significant inbreeding depression in caribou through increased juvenile mortality, Gagnon et al. (2019) detected no clear fitness effects in certain wild caribou populations. However, the negative impacts of inbreeding on survival and reproductive success are well-documented across diverse mammals, including other ungulates like soay sheep (*Ovis aries*) and red deer (B ér é nos et al., 2016; Hewett et al., 2024), suggesting that the high F_ROH_ in declining southern ranges warrants further research. This includes assessing whether inbreeding depression is amplified under changing environmental conditions (Colpitts et al., 2024, Hewett et al., 2025). Notably, elevated inbreeding was not restricted to small populations; with larger groups such as Itcha-Ilgachuz also showing high F_ROH_. This likely reflects a legacy of historical isolation and genetic drift rather than recent demographic decline alone (Deakin et al., 2025; Taylor et al., 2020, 2024). Such prolonged isolation and associated genetic drift and small N_e_ would facilitate the accumulation of ROH (Colpitts et al., 2022), regardless of N_c._ This underscores that N_c_ is an insufficient proxy for genetic risk, particularly in evolutionarily unique lineages where historical isolation has shaped current genomic diversity.

Most subpopulations in the Central-eastern, Jasper-Banff, and South-eastern metapopulations exhibited contemporary N_e_ point estimates below the critical threshold of N_e_ = 50, indicating high short-term inbreeding risk. This concern is likely broader than estimates alone suggest, as N_e_ is probably also below 50 in many subpopulations for which estimation was not possible due to very low N_c_ and insufficient sample size. While most subpopulations had N_e_ point estimates below 500 (the threshold associated with long-term maintenance of adaptive capacity; Jamieson & Allendorf, 2012), only 22 of 32 had 90% CIs situated entirely below this value. For subpopulations with CIs overlapping these thresholds, uncertainty in N_e_ estimates complicates conservation prioritization (reviewed in Clarke et al., 2024), highlighting the need for either increased genetic sampling to improve estimate precision or precautionary management approaches that treat uncertain subpopulations with the same urgency as those with confirmed low N_e_ (Waples, 2025). In contrast, northern subpopulations, particularly Graham, Finlay, and Pink Mountain, maintained substantially higher N_e_ values, with Graham being the only subpopulation whose point estimate exceeded 500 with CIs largely above this threshold, suggesting better genetic health and lower immediate extinction risk in this region.

While metapopulation N_e_ can theoretically exceed the sum of subpopulation N_e_ under complete isolation, our findings indicate that caribou metapopulations exhibit reduced N_e_ (<500) relative to expectations, likely resulting from genetic drift, asymmetric gene flow, and unequal subpopulation contributions (Caballero, 2020; Gomez-Uchida et al., 2013; Kurland et al., 2023). Although, these metapopulation estimates should be interpreted cautiously, as both currentNe2 and GONE2 assume panmixia within subpopulations, an assumption that can lead to underestimation when gene flow inflates LD patterns (Santiago et al., 2025). While structure correction methods exist, they require assumptions about migration symmetry and equal subpopulation sizes that caribou violate (Government of British Columbia, 2025; Theoret et al., 2022), making our estimates likely on the lower bounds. The observed variation in metapopulation N_e_ appears to be linked to spatial structure and connectivity (Deakin et al., 2025). In particular, the South-eastern and Jasper-Banff metapopulations, characterized by linear connectivity along the Rocky Mountains (Cavedon et al., 2023; Deakin et al., 2025), exhibit the lowest N_e_ (< 60). Such linear stepping-stone arrangements are particularly vulnerable: loss of critical central subpopulations like Columbia North could sever gene flow and trigger cascading metapopulation collapse (Kurland et al., 2023). In contrast, North-western, North-eastern, and Central-eastern metapopulations possibly maintain higher N_e_ (> 150) through more complex spatial arrangements providing greater connectivity redundancy (Cichowski & Ray, 2022; Deakin et al., 2025). These patterns likely reflect differences in migratory behavior and landscape connectivity among regions, which together shape opportunities for gene flow among subpopulations (Lamb et al., 2025; Theoret et al., 2022). Northern subpopulations (Graham, Finlay, and Pink Mountain) may serve as genetic reservoirs, exhibiting elevated diversity possibly linked to admixture between ancestral lineages and known migration corridors (Deakin et al., 2025; Hughes et al., 2025; Leech et al., 2016), though how movement patterns translate into effective gene flow and whether habitat fragmentation is disrupting historical connectivity warrants further investigation.

N_e_/N_c_ ratios averaged a median of 0.462 across subpopulations with *n* > 10, exceeding values reported for other polygynous ungulate species (Frankham, 1995), such as moose (*Alces alces*, 0.28; Lee et al., 2020), bighorn sheep (*Ovis canadensis*, 0.33; FitzSimmons et al., 1995), and European reindeer (0.124; Kvalnes et al., 2024), as well as the typical mammalian ratio of 0.3 (Frankham, 2021; Hoban et al., 2024). However, lower ratios were observed in several Central-eastern subpopulations (*e.g.*, Kennedy-Siding: 0.213, Klinse-Za: 0.355, A-La-Peche: 0.259) and individual subpopulations in other metapopulations (Carcross: 0.029, Gataga: 0.134, Columbia North: 0.196), with only Carcross and Gataga falling below the conventional threshold of 0.1, indicative of potential genetic vulnerability (Frankham, 1995; Hoban et al., 2020). Smaller N_e_/N_c_ ratios may result from strong male-male competition for mates, leading to high variance in reproductive success (McFarlane et al., 2018), which could be exacerbated by local population dynamics that concentrate breeding opportunities among fewer individuals (McCullough, 1999; Wittmer et al., 2010). Habitat fragmentation and limited gene flow may also contribute to low ratios, particularly in isolated or declining subpopulations (Environment Canada, 2014). Seven subpopulations exhibited N_e_/N_c_ ratios exceeding 1 (Telkwa, Brazeau, Maxhamish, Little-Rancheria, Tsenaglode, Central Selkirks, Graham), contrary to the expectation that N_e_ is typically lower than N_c_ in wild populations (Waples 2024b). Possible explanations include temporal incongruence between N_e_ and N_c_ estimates, variation in census techniques (Government of British Columbia, 2025), uncertainty in N_c_ estimates, limited genetic sampling, and undetected migration between subpopulations (Hedrick 2005; Hedgecock, 1992). We recommend interpreting N_e_/N_c_ cautiously at local scales, particularly where genetic and census sampling periods do not overlap.

Mean F_ROH_ was negatively associated with N_c_ but not with contemporary N_e_. Results for N_c_ align with previous findings in caribou subpopulations (Serrouya et al., 2012) and across various wild species with limited population sizes (Åkesson et al., 2016; Peart et al., 2020). This pattern reflects how reduced population size increases genetic drift and decreases gene flow, thereby elevating homozygosity (Kardos et al. 2018). The quadratic census model revealed a non-linear N_c_-F_ROH_ relationship, with inbreeding effects appearing to plateau at larger population sizes, possibly reflecting limited gene flow even among large but fragmented populations. In contrast, N_e_ showed weak predictive power, indicating that recent demographic changes may be the primary driver of current inbreeding patterns rather than long-term effective breeding dynamics (Kardos et al. 2018).

Temporal N_e_ analyses suggested multiple waves of population decline across subpopulations, broadly progressing from north to south over several centuries. Results from GONE should be interpreted as general trends rather than precise timing, given inherent uncertainty in recombination rate and generation time assumptions, and because simulation studies have shown that GONE may not perfectly track the magnitude or timing of N_e_ changes (Santiago et al., 2020). Using an 8-year generation time, the earliest signal was detected in the North-eastern metapopulation (∼1400, based on two subpopulations and metapopulation-level results), tentatively coinciding with the onset of the Little Ice Age (LIA; c. 1350 in North America; Luckman et al., 2020). Increased winter severity and icing events associated with this period may have reduced forage access and survival, contributing to demographic pressure, though climatic effects should be interpreted as contributing factors rather than sole causal drivers (Bergerud, 1974; Vors & Boyce, 2009). Subsequent declines were detected first in northern subpopulations and Itcha-Ilgachuz (40-30 generations ago, ∼1700-1780), followed by central subpopulations (30-20 generations ago, ∼1780-1860) and southern subpopulations (20-10 generations ago, ∼1860-1940), broadly overlapping with the later stages and end of the LIA (c. 1850; Luckman et al., 2020), which brought rapid ecological changes including increased fire frequency, glacier retreat, and shifts in snow-crusting patterns that impeded movement and predator avoidance (Harvey & Smith, 2017; Starheim et al., 2013; Wood & Smith, 2013). Declines in southern and central regions were likely further compounded during the settlement era (∼1860 onwards) by anthropogenic pressures including the gold rush, railway expansion (1885), and market hunting (Mackie, 2020; McLellan, 2010; Spalding, 2000). Recombination rate and generation time variation introduce modest temporal uncertainty (±10-20 years), yet the relative north-to-south gradient and the overlap with known environmental and anthropogenic events remain clear.

More recent bottlenecks appear to have occurred in distinct waves, largely tracking the historical progression of land-use changes. Bottlenecks occurred 20–30 generations ago (∼1850–1900) in Central-eastern and Jasper-Banff metapopulations and extended into the last ten generations (∼1950) in South-eastern subpopulations, coinciding with the peak of Euro-American settlement. This period brought intensified anthropogenic pressures through logging, mining, hunting, and gold rushes (*e.g*., Fraser Canyon 1858, Cariboo 1860s; reviewed in Mackie, 2020). British Columbia became a colony in 1858, joined Confederation in 1871, and was connected by rail in 1885, fueling industrial expansion (McGillivray, 2011; McLellan, 2010; Santomauro et al., 2012; Sutherland, 2014). Hunting became a major factor in the decline of southern caribou during the 19th century, leading to sex-specific regulations beginning in the 1890s and progressive hunting closures across British Columbia, starting in 1914 and expanding throughout the 20th century (see Spalding, 2000). Moose and wolves expanded into central British Columbia in the early 20th century, increasing predation pressure (Bergerud & Elliot, 1986; Seip & Cichowski, 1996). In Jasper-Banff specifically, extensive wolf control programs (∼1900–1959) followed this settlement-era bottleneck, enabling demographic recovery that peaked around 1965 (Bradley & Neufeld, 2012). These settlement-period impacts produced genetic signatures evident in Central-eastern, Jasper-Banff, and South-eastern metapopulations (Laliberte & Ripple, 2004).

While the earlier declines reflect climatic and historical anthropogenic pressures, N_e_ declines over the last 5-10 generations (∼1940–1980) evident at the metapopulation scale point to intensifying isolation driven by industrial-era land-use change, though model assumptions regarding population structure may also contribute to these signals (Santiago et al., 2025). The abundance of long ROH segments provides independent evidence of close-relative mating within the last few generations, consistent with these reconstruction results. Although caribou numbers dropped 50-60% since the 1970s (Ministry of Forests, 2018), this rapid crash was likely not fully captured in genetic reconstructions due to the persistence of ancestral polymorphism. From the 1940s onward, industrial logging, hydroelectric development, road construction, oil and gas exploration, and recreational expansion caused widespread habitat disturbance across British Columbia (Edwards, 1954; Hebblewhite, 2017; Lochhead et al., 2022; Konkolics et al., 2021; Wilson et al., 2022), triggering apparent competition dynamics whereby habitat modification favored moose and deer expansion and consequently increased wolf predation on caribou (Mumma et al., 2018; Neufeld et al., 2021). Jasper-Banff represents a notable exception, with genetic reconstructions showing a recovery signal during this period, likely reflecting the demographic recovery described above (Bradley & Neufeld, 2012).

Small increases or stabilization in N_e_ over the last five to ten generations (∼1960 to 1990) were apparent in several northern subpopulations. These trends are consistent with the timing of various management interventions including long-standing male-only hunting restrictions (implemented intermittently since 1893 and more widely by the 1970s) and extensive predator control programs (∼1900 onwards), which have both been associated with population size increases across western Canada (Bergerud & Elliot, 1986; McLellan, 2010; Spalding, 2000, Lamb et al., 2022). However, these trends are not reflected at the metapopulation level. This discrepancy could be explained by several factors. First, habitat loss and fragmentation across western Canada over the past 30 years (Dickie et al., 2023; Lochhead et al., 2022; Wittmer et al., 2007) may be limiting gene flow between recovering local subpopulations, preventing management effects across the broader metapopulation. Second, while some subpopulations show minor improvements, others may still be experiencing continued decline, effectively cancelling out genetic gains at the metapopulation scale (Lamb et al., 2024). Third, the time lag between local conservation interventions and detectable genetic impacts at the metapopulation level may be substantial, potentially requiring several more generations before changes N_e_ become apparent (Shaw et al., 2025). This emphasizes the fact that short-term management actions might not be detected genetically if not conducted over multiple generations (Nagy-Reis et al., 2021; Palm et al., 2020). Fourth, this discrepancy may also stem from methodological limitations, as the signature of localized demographic growth can be masked by broader signals of genetic isolation in structured populations, potentially leading models to underrepresent the efficacy of recent management actions (Santiago et al., 2025).

### Conservation implications and recommendations

The low N_e_ and elevated F_ROH_ values observed across several subpopulations and metapopulations suggest that genetic erosion is compounding the demographic threats already imposed by habitat loss and predation. Populations with N_e_ < 50 face elevated extinction risk from inbreeding depression, while those with N_e_ < 500 may face reduced adaptive potential; combined with F_ROH_ > 0.1, these thresholds suggest that some subpopulations could have limited recovery potential even if habitat is restored, pointing to the utility of genetic thresholds, alongside connectivity and status assessments, as monitoring tools for long-term viability (Government of British Columbia, 2025; Kyriazis et al., 2020; Robinson et al., 2023). These findings underscore the value of integrating genomic metrics into caribou recovery frameworks to identify subpopulations where demographic and genetic pressures converge.

Where inbreeding is severe, genetic rescue could offer benefits (Poirier et al., 2019; Sundell et al., 2023), though past translocation efforts in this system have shown limited success (Boutin and Merrill, 2016; Leech et al., 2017; Young et al., 2001), and risks such as disruption of learned movement patterns, pathogen introduction, and outbreeding depression warrant consideration (Gerhold and Hickling, 2016; Mellya et al., 2023). When translocations are pursued, screening for pathogens, genetic load and behavioral compatibility would help mitigate these risks (Taylor et al., 2024; Hughes et al., 2025; Sherman et al., 2025).

Demographic and habitat-based strategies, including predator management, maternal penning, and restoration of landscape connectivity, remain central to caribou recovery and should naturally increase Ne and facilitate gene flow, particularly across northern metapopulations (Dickie et al., 2023; Grant et al., 2019; Hervieux et al., 2014; Lamb et al., 2024; Leech et al., 2016; McLellan et al., 2025; Serrouya et al., 2019). Sustained genomic monitoring, ideally developed in partnership with First Nations (Lamb et al., 2022), would provide the evidence needed to track subpopulation viability over time and ensure recovery efforts address both immediate demographic pressures and longer-term genetic resilience.

## Acknowledgements

We gratefully acknowledge the Government of British Columbia, students, Parks Canada partners, First Nations communities, and all other collaborators who contributed to the collection of samples used in this study. We acknowledge that this research took place on the traditional territories of First Nations and M étis peoples, where caribou persist today and would not have been possible without their collaboration. This work was financially supported by the Government of British Columbia, Environment and Climate Change Canada’s Canadian Wildlife Service, Parks Canada, and Natural Sciences and Engineering Research Council of Canada (NSERC) grants to Jocelyn Poissant and Marco Musiani.

## Author Contributions

**Charlotte Bourbon**: Conceptualization, Methodology, Formal Analysis, Investigation, Data Curation, Writing – Original Draft, Review & Editing, Visualization, Project Administration. **Samuel Deakin**: Methodology, Data curation, Lab work, Review & Editing. **Anita Michalak**: Methodology, Data curation, Lab work, Review & Editing. **Margaret Hughes**: Data curation, Visualization, Review & Editing, Investigation. **Maria Cavedon**: Data Curation, Review & Editing. **Lalenia Neufeld**: Conceptualization, Data curation, Methodology, Review & Editing, Investigation. **Agnes Pelletier**: Conceptualization, Data Curation, Review & Editing. **Jean Polfus**: Conceptualization, Review & Editing, Funding Acquisition. **Helen Schwantje**: Conceptualization, Methodology, Data Curation, Review & Editing, Funding Acquisition. **Caeley Thacker**: Conceptualization, Methodology, Review & Editing, Project Administration, Funding Acquisition. **Marco Musiani**: Conceptualization, Supervision, Writing – Review & Editing, Project Administration, Funding Acquisition. **Jocelyn Poissant**: Conceptualization, Methodology, Supervision, Writing – Review & Editing, Project Administration, Funding Acquisition.

