## Supporting Information for "Inbreeding and demographic history of caribou (*Rangifer tarandus*) in western Canada inferred from genome-wide SNP data"

#### 1. Parameters comparison for detecting ROHs

While parameters can strongly impact the number and length of ROHs, and therefore $F_{ROH}$ (Ceballos et al. 2018), there is little agreement on optimal parameter values in the literature (Peripolli et al. 2017). This lacks of consensus partly reflects differences in SNP density across datasets, which directly affects appropriate parameterization (Ferencakovic et al. 2011; Meyermans et al. 2020). We thus first compared results obtained when using different combinations of parameters. Specifically, the minimum number of SNPs within a ROH (*minSNP*) was allowed to be either 20, 30, 40, or 50, the maximum number of heterozygous SNPs (*maxOppRun*) was allowed to vary between 0 and 4, and the maximum number of missing genotypes in a run (*maxMissRun*) was allowed to vary from 0 to 2. The minimum length of a run was set at 300 kb, and the maximum gap between consecutive SNPs was set to 1 Mb, as suggested by Biscarini et al. (2018).

To evaluate the impacts of parameter choices and select a combination for our ROH detection, we fit linear models with the minimum number of SNPs within a ROH, the maximum number of heterozygous SNPs, and the maximum number of missing genotypes in a run as categorical predictors, including two-way interactions. Using Type III ANOVA, we determined which parameters significantly affected ROH estimation, while partial $\eta^2$ values quantified their relative importance and effect sizes. These effect sizes ($\eta^2$) directly measure how much variance in $F_{ROH}$ and ROH counts is explained by each parameter, with values above 0.14 considered large effects (Richardson 2011).

Our analysis of parameter effects on ROH estimation revealed that *minSNP* had the strongest influence on both $F_{ROH}$ and ROH count (partial $\eta^2$ = 0.15 and 0.77, respectively; both $p < 0.001$), followed by the interaction between *minSNP* and *maxOppRun* (partial $\eta^2$ = 0.10 and 0.69, $p < 0.001$), and *maxOppRun* (partial $\eta^2$ = 0.08 and 0.52, $p < 0.001$). In particular, the effect of increasing *maxOppRun* is substantially more pronounced at lower *minSNP* thresholds (particularly between 20 and 30), where permitting more heterozygous SNPs dramatically increases both $F_{ROH}$ values (Figure S1 A) and ROH counts (Figure S1 B), suggesting that parameter selection is especially critical when using less stringent SNP density requirements.

***minSNP* parameter**: For *minSNP*, we selected a threshold of 20 to balance sensitivity and specificity. This choice relates to our medium-density SNP array characteristics (1 SNP every ~60 kb on average; Carrier et al., 2022). With a genome size estimated by detectRUNS to be approximately 2 Gb (based on the 33,346 SNPs used in our analyses), the smallest ROH theoretically detectable with *minSNP* = 20 should be ~1.2 Mb in length (20 SNPs × 60 kb average distance). This means our analysis primarily captures ROH segments >1.2 Mb to longer segments (>8 Mb), which typically reflect inbreeding events from recent to intermediate timeframes rather than very distant ancestral relatedness.

***maxOppRun* parameter:** For *maxOppRun*, we selected a value of 2 after comparing its effects across different *minSNP* thresholds. This parameter showed significant influence on ROH detection (partial $\eta^2 = 0.08$ and 0.52 for $F_{ROH}$ and ROH count, respectively). While *maxOppRun* = 4 produced substantially higher $F_{ROH}$ values and ROH counts (especially at *minSNP* = 20), suggesting potential inflation of ROH detection by including segments with excessive heterozygosity, values of 0-1 appeared overly stringent, potentially missing genuine ROH. At our selected *minSNP* = 20, *maxOppRun* = 2 appears to tolerate a biologically reasonable number of heterozygous SNPs while maintaining specificity against false positives.

**maxMissRun parameter:** The *maxMissRun* parameter exhibited minimal impact on ROH estimation (partial $\eta^2 < 0.001$, $p < 0.05$). Although it demonstrated negligible influence, we set this parameter to 2 to allow for a reasonable tolerance of missing data in our analysis.

Our final parameter selection (*minSNP* = 20, *maxOpprun* = 2, *maxMissRun* = 2) prioritizes the detection of high-confidence homozygous regions. By using a *minSNP* of 20, we ensure that detected ROHs are statistically unlikely to arise by chance, given our array density. Based on an average marker spacing of ~60 kb, this configuration is optimized to detect segments ≥ 1.20 Mb. While the flexibility of these parameters (allowing 2 heterozygous or missing calls) could theoretically allow the detection of shorter segments in SNP-dense regions or at chromosome ends, we interpret results in the 'Short' category (specifically < 1.20 Mb) with caution, as they represent the lower limit of our array 's resolution.

### 2. Demographic scenarios with DIYABC-RF

Subpopulation datasets were filtered for missing data using BCFtools v1.17 (SAMtools v1.21), chosen over PLINK for faster processing. Only polymorphic sites were retained, and datasets were converted using 'vcf2DIABC.py' script (https://github.com/loire/vcf2DIYABC.py) in Python v3.12.5. DIYABC-RF employs a two-step process: first simulating genetic datasets under various $N_e$ trajectories, then using RF classification to compare summary statistics between observed and simulated data, allowing robust discrimination between competing demographic scenarios without requiring explicit likelihood calculations (Collin et al. 2021). A sex ratio of 58% females to 42% males (NM = 1.38F) was assumed based on Wittmer et al. (2010) who conducted multi-year monitoring of radio-collared caribou across British Columbia. All four scenarios received equal prior probabilities (0.25). $N_e$ priors (10-1,000 for contemporary N0 and 10-10,000 for historical N1 and N2) were calibrated based on extensive literature review and preliminary data exploration. The narrower range for contemporary $N_e$ was informed by recent population estimates and genetic studies suggesting current caribou population sizes are generally smaller than historical levels (Weckworth et al. 2018; Yannic et al. 2014), while the broader ranges for N1 and N2 allowed for exploration of various historical population sizes given the uncertainty in pre-decline demographics. Time priors were set at t1 (1-5 generations) for recent demographic events and t2 (25-100 generations) for historical events, representing Bayesian prior distributions from which the ABC framework samples during simulation to estimate posterior distributions of event timing. Assuming a generation time of 8 years (Government of British Columbia, 2025), this translates to events occurring ~8-40 years ago and ~200-800 years ago, corresponding to recent bottlenecks likely due to anthropogenic habitat alteration (Spalding 2000) and potential historical bottlenecks. For each scenario, we ran 20,000 simulations (80,000 total), with random forest analysis using 10 noise variables and 2,000 trees to balance precision and computational efficiency (Collin et al. 2021).

### Supplementary Tables and Figures

*Table S1: One-sample Wilcoxon signed-rank test results for comparing $F_{ROH}$ inbreeding coefficients of each subpopulation to the overall median.* The table shows test statistics and p-values for one-sided tests (greater and less than the overall median). Five subpopulations had sample sizes smaller than 3 and were not compatible with one-sample Wilcoxon signed-rank tests.

| Subpopulation | W-statistic | p-value (Greater) | p-value (Less) |
|---|---|---|---|
| Columbia-North | 1225 | 1.78E-15 | 1 |
| Itcha-Ilgachuz | 3003 | 1.26E-14 | 1 |
| Tweedsmuir | 435 | 1.86E-09 | 1 |
| Tonquin | 401 | 3.73E-08 | 1 |
| Maligne | 55 | 0.000977 | 1 |
| Chinchaga | 199 | 0.0086 | 0.992 |
| Wells-Gray-South | 151 | 0.0115 | 0.990 |
| Brazeau | 21 | 0.0156 | 1 |
| Central-Selkirks | 21 | 0.0156 | 1 |
| Barkerville | 15 | 0.0312 | 1 |
| Groundhog | 20 | 0.0313 | 0.984 |
| Carcross | 18 | 0.0781 | 0.953 |
| Atlin | 128 | 0.0978 | 0.909 |
| A-La-Peche | 116 | 0.0982 | 0.909 |
| Purcells-South | 6 | 0.125 | 1 |
| Westside-Fort-Nelson | 16 | 0.406 | 0.656 |
| Telkwa | 9 | 0.406 | 0.688 |
| Quintette | 103 | 0.536 | 0.478 |
| Wolverine | 436 | 0.578 | 0.426 |
| Klinse-Za (Moberly) | 291 | 0.748 | 0.257 |
| East-Wiliston | 2 | 0.75 | 0.375 |
| Calendar | 8 | 0.963 | 0.0488 |
| Maxhamish | 4 | 0.99 | 0.0137 |

| | | | |
|---|---|---|---|
| Gataga | 1 | 0.984 | 0.0313 |
| Graham | 55 | 0.991 | 0.00948 |
| Kennedy-Siding | 20 | 0.991 | 0.0108 |
| North-Cariboo | 60 | 0.996 | 0.00436 |
| Snake-Sahtaneh | 37 | 0.998 | 0.00244 |
| Hart-Ranges | 360 | 1 | 9.72E-07 |
| Chase | 54 | 1 | 1.45E-09 |
| Pink-Mountain | 127 | 1 | 3.13E-05 |
| Muskwa | 67 | 1 | 0.000334 |
| Redrock-Prairie-Creek | 0 | 1 | 0.000488 |
| Little-Rancheria | 0 | 1 | 0.00781 |
| Tsenaglode | 0 | 1 | 0.00781 |
| Frog | 0 | 1 | 0.00781 |
| Finlay | 0 | 1 | 0.0312 |
| Horseranch | 0 | 1 | 0.0625 |
| Level-Kawdy | 0 | 1 | 0.125 |
| Narraway | 0 | 1 | 0.125 |

***Table S2. Contemporary effective population size ($N_e$) estimates for 31 caribou subpopulations at recombination rates of 0.9 and 1.1 cM/Mb.*** Subpopulations with small sample sizes (n ≤ 5) or $N_c$ = 0 (Maligne) were excluded from analyses.

| **Subpopulation** | **n** | **r = 0.9 cM/Mb** | | | **r = 1.1 cM/Mb** | | |
|---|---|---|---|---|---|---|---|
| | | $N_e$ | 90% CI | $N_e/N_c$ (90% CI) | $N_e$ | 90% CI | $N_e/N_c$ (90% CI) |
| A-La-Peche | 18 | 35.38 | 27.32–45.80 | 0.236 (0.182–0.305) | 34.57 | 26.36–45.35 | 0.230 (0.176–0.302) |
| Atlin | 19 | 1074.19 | 427.43–2699.56 | 0.703 (0.280–1.768) | 1015.40 | 391.23–2635.37 | 0.665 (0.256–1.726) |
| Banff | 2 | NA | NA | NA | NA | NA | NA |
| Barkerville | 5 | 15.60 | 8.80–27.65 | 0.312 (0.176–0.553) | 15.20 | 8.34–27.69 | 0.304 (0.167–0.554) |
| Brazeau | 6 | 24.12 | 13.52–43.04 | 2.412 (1.352–4.304) | 23.55 | 12.83–43.22 | 2.355 (1.283–4.322) |
| Calendar | 9 | 105.58 | 51.59–216.07 | 0.457 (0.223–0.935) | 101.90 | 48.26–215.14 | 0.441 (0.209–0.931) |
| Carcross | 6 | 25.45 | 14.10–45.94 | 0.030 (0.017–0.054) | 24.89 | 13.39–46.25 | 0.029 (0.016–0.054) |
| Central-Selkirks | 6 | 213.57 | 57.56–792.63 | 7.910 (2.132–29.357) | 204.60 | 52.45–798.08 | 7.578 (1.943–29.559) |
| Chase | 45 | 219.96 | 173.34–279.11 | 0.367 (0.289–0.465) | 210.78 | 164.45–270.16 | 0.351 (0.274–0.450) |
| Chinchaga | 22 | 192.81 | 126.63–293.58 | 0.599 (0.393–0.912) | 185.01 | 119.38–286.74 | 0.575 (0.371–0.891) |
| Columbia-North | 49 | 43.63 | 38.86–49.00 | 0.199 (0.177–0.224) | 42.42 | 37.56–47.91 | 0.194 (0.172–0.219) |
| Columbia-South | 2 | 12.40 | 5.01–39.15 | 12.40 (5.01–39.15) | 12.30 | 5.01–41.23 | 12.30 (5.01–41.23) |

| | | | | | | | |
|---|---|---|---|---|---|---|---|
| East-Williston | 3 | 28.54 | 9.45–86.21 | 0.182 (0.100–0.330) | 27.88 | 8.77–88.63 | 0.177 (0.095–0.332) |
| Finlay | 5 | NA | NA | 0.135 (0.076–0.241) | NA | NA | 0.133 (0.072–0.243) |
| Frog | 7 | 37.39 | 20.59–67.90 | 12.447 (4.844–31.981) | 36.54 | 19.54–68.33 | 11.731 (4.423–31.118) |
| Gataga | 6 | 24.24 | 13.61–43.17 | 0.190 (0.128–0.281) | 23.72 | 12.94–43.47 | 0.187 (0.124–0.282) |
| Graham | 22 | 1593.20 | 620.07–4093.53 | 0.537 (0.445–0.647) | 1501.60 | 566.10–3983.04 | 0.513 (0.422–0.624) |
| Groundhog | 6 | 8.74 | 5.91–12.93 | 0.555 (0.467–0.658) | 8.58 | 5.69–12.95 | 0.530 (0.443–0.633) |
| Hart-Ranges | 66 | 281.72 | 233.48–339.92 | 0.215 (0.162–0.285) | 269.33 | 221.48–327.52 | 0.210 (0.156–0.282) |
| Hay-River | 1 | NA | NA | 0.361 (0.309–0.423) | NA | NA | 0.350 (0.297–0.412) |
| Horseranch | 4 | 962.40 | 59.89–15466.18 | 1.227 (0.180–8.353) | 910.40 | 53.15–15587.49 | 1.161 (0.161–8.373) |
| Itcha-Ilgachuz | 77 | 310.09 | 261.34–367.93 | 2.131 (0.804–5.652) | 296.10 | 247.79–353.83 | 2.041 (0.741–5.624) |
| Kennedy-Siding | 15 | 29.64 | 22.34–39.31 | 0.378 (0.249–0.573) | 28.94 | 21.52–38.94 | 0.361 (0.234–0.556) |
| Klinse-Za | 37 | 49.85 | 42.59–58.34 | 1.123 (0.762–1.655) | 48.24 | 40.92–56.88 | 1.078 (0.720–1.615) |
| Level-Kawdy | 3 | 933.41 | 38.38–22702.00 | 0.890 (0.623–1.271) | 883.20 | 34.23–22803.36 | 0.847 (0.585–1.227) |
| Little-Rancheria | 7 | 923.02 | 135.63–6281.29 | 0.480 (0.355–0.650) | 873.43 | 121.16–6296.62 | 0.466 (0.339–0.640) |
| Maligne | 10 | 38.19 | 24.51–59.51 | 0.459 | 37.35 | 23.45–59.51 | 0.446 |

| | | | | | | | |
|---|---|---|---|---|---|---|---|
| | | | | (0.298–0.706) | | | (0.284–0.701) |
| Maxhamish | 9 | 238.72 | 90.05–632.97 | 0.753 (0.469–1.210) | 228.57 | 82.94–629.89 | 0.722 (0.440–1.182) |
| Muskwa | 29 | 346.51 | 228.51–525.43 | 0.989 (0.809–1.209) | 330.87 | 214.59–510.15 | 0.962 (0.779–1.187) |
| Narraway | 3 | 839.43 | 36.91–19089.56 | 1.915 (0.197–18.637) | 795.04 | 32.89–19215.56 | 1.804 (0.175–18.646) |
| Narrow-Lake | 2 | 599.36 | 18.06–19888.10 | 0.900 (0.661–1.226) | 569.09 | 16.24–19939.15 | 0.865 (0.626–1.194) |
| North-Cariboo | 24 | 189.81 | 128.81–279.70 | 0.419 (0.299–0.587) | 182.22 | 121.64–272.97 | 0.406 (0.286–0.578) |
| Pink-Mountain | 40 | 474.33 | 332.08–677.50 | 1.020 (0.327–3.182) | 451.55 | 311.74–654.06 | 0.977 (0.299–3.189) |
| Purcells-South | 3 | 12.80 | 5.55–29.52 | 0.612 (0.479–0.782) | 12.80 | 5.29–30.97 | 0.587 (0.455–0.758) |
| Quintette | 20 | 67.20 | 49.64–90.98 | 0.236 (0.182–0.305) | 65.20 | 47.48–89.54 | 0.230 (0.176–0.302) |
| Redrock-Prairie-Creek | 11 | 44.04 | 28.62–67.77 | 0.703 (0.280–1.768) | 42.82 | 27.26–67.27 | 0.665 (0.256–1.726) |
| Snake-Sahtaneh | 21 | 236.56 | 147.23–380.09 | NA | 226.57 | 138.27–371.26 | NA |
| South-Selkirks | 2 | NA | NA | 0.312 (0.176–0.553) | NA | NA | 0.304 (0.167–0.554) |
| Telkwa | 5 | 83.20 | 28.37–243.98 | 2.412 (1.352–4.304) | 80.40 | 26.19–246.84 | 2.355 (1.283–4.322) |
| Tonquin | 28 | 49.45 | 40.46–60.43 | 0.457 (0.223–0.935) | 48.08 | 38.95–59.33 | 0.441 (0.209–0.931) |
| Tsenaglode | 7 | 1608.40 | 165.25–15654.97 | 0.030 (0.017–0.054) | 1515.70 | 146.73–15662.98 | 0.029 (0.016–0.054) |

| | | | | | | | |
|---|---|---|---|---|---|---|---|
| Tweedsmuir | 29 | 160.24 | 117.62–218.30 | 7.910 (2.132–29.357) | 153.94 | 111.51–212.51 | 7.578 (1.943–29.559) |
| Wells-Gray-South | 19 | 78.40 | 55.98–109.79 | 0.367 (0.289–0.465) | 76.00 | 53.43–108.11 | 0.351 (0.274–0.450) |
| Westside-Fort-Nelson | 7 | 206.99 | 66.33–645.91 | 0.599 (0.393–0.912) | 198.40 | 60.79–647.37 | 0.575 (0.371–0.891) |
| Wolverine | 42 | 202.00 | 158.13–258.03 | 0.199 (0.177–0.224) | 193.81 | 150.14–250.17 | 0.194 (0.172–0.219) |

97

***Table S3. Contemporary effective population size ($N_e$) estimates for caribou metapopulations at recombination rates of 0.9 and 1.1 cM/Mb.*** All the samples (*n* = 759) were used for this analysis.

| **Metapopulation** | **n** | **r = 0.9 cM/Mb** | | **r = 1.1 cM/Mb** | |
|---|---|---|---|---|---|
| | | $N_e$ | 90% CI | $N_e$ | 90% CI |
| Central-eastern | 220 | 317.36 | 297.26–338.83 | 303.30 | 283.21–324.80 |
| Itcha-Ilgachuz | 77 | 310.09 | 261.34–367.93 | 296.10 | 247.79–353.83 |
| Jasper-Banff | 46 | 45.43 | 40.09–51.47 | 44.22 | 38.79–50.42 |
| North-eastern | 181 | 449.95 | 410.52–493.16 | 428.37 | 389.34–471.32 |
| North-western | 167 | 155.04 | 145.65–165.04 | 149.05 | 139.59–159.16 |
| South-eastern | 68 | 59.20 | 53.71–65.26 | 57.31 | 51.75–63.48 |

*Table S4: Model comparison results for assessing the relationship between inbreeding coefficients $F_{ROH}$ and log-transformed population metrics ($N_c$, $N_e$) and their interactions.* Each model is evaluated based on the following metrics: AICc: the corrected Akaike Information Criterion; $R^2$: the coefficient of determination, representing the proportion of variance explained by the model; ΔAICc: the difference in AICc between the model and the best-fitting model; and Weight: The relative likelihood of the model, given the set of models compared. Only models with ΔAICc < 2 were retained for model averaging. Subpopulations with small sample sizes ($n \leq 5$) or $N_c = 0$ (Maligne) were excluded from analyses.

| Model | AICc | $R^2$ | ΔAICc | Weight |
|---|---|---|---|---|
| $\log(N_c)$ | 33.062 | 0.19 | 0 | 0.426 |
| $\log(N_c)+ \log(N_c)^2$ | 33.275 | 0.252 | 0.213 | 0.383 |
| $\log(N_c) + \log(N_e)$ | 35.693 | 0.191 | 2.631 | 0.114 |
| $\log(N_e)$ | 38.044 | 0.049 | 4.983 | 0.035 |
| $\log(N_c) + \log(N_e)$ + interaction | 38.554 | 0.191 | 5.492 | 0.027 |
| $\log(N_e) + \log(N_e)^2$ | 40.02 | 0.07 | 6.958 | 0.013 |

***Table S5: Model-averaged parameter estimates for the top models (ΔAICc < 2) explaining the relationship between inbreeding coefficients $F_{ROH}$ and log-transformed population census size ($N_c$).*** Model averaging was performed across the simple linear model (log(Nc)) and the quadratic model ($\log(N_c) + \log(N_c)^2$). Parameter estimates for individual models and model-averaged estimates (conditional averages) are shown. The quadratic term appears only in Model 2 (importance = 0.46). Subpopulations with small sample sizes ($n \leq 5$) or $N_c = 0$ (Maligne) were excluded from analyses.

| **Model** | **Parameter** | **Estimate** | **Std. Error** | **t/z value** | **Pr(>\|t/z\|)** | **$R^2$** |
|---|---|---|---|---|---|---|
| Model-averaged | Intercept | -0.977 | 1.039 | 0.941 | 0.347 | NA |
| | $\log(N_c)$ | -0.447 | 0.413 | 1.081 | 0.28 | NA |
| | $\log(N_c^2)$ | 0.059 | 0.041 | 1.448 | 0.148 | NA |
| Model 1: $\log(N_c)$ | Intercept | -1.664 | 0.366 | -4.55 | <0.001 | 0.19 |
| | $\log(N_c)$ | -0.173 | 0.067 | -2.586 | 0.015 | 0.19 |
| Model 2: $\log(N_c)+\log(N_c^2)$ | Intercept | -0.203 | 1.043 | -0.195 | 0.847 | 0.252 |
| | $\log(N_c)$ | -0.784 | 0.415 | -1.888 | 0.069 | 0.252 |
| | $\log(N_c^2)$ | 0.059 | 0.041 | 1.448 | 0.148 | 0.252 |

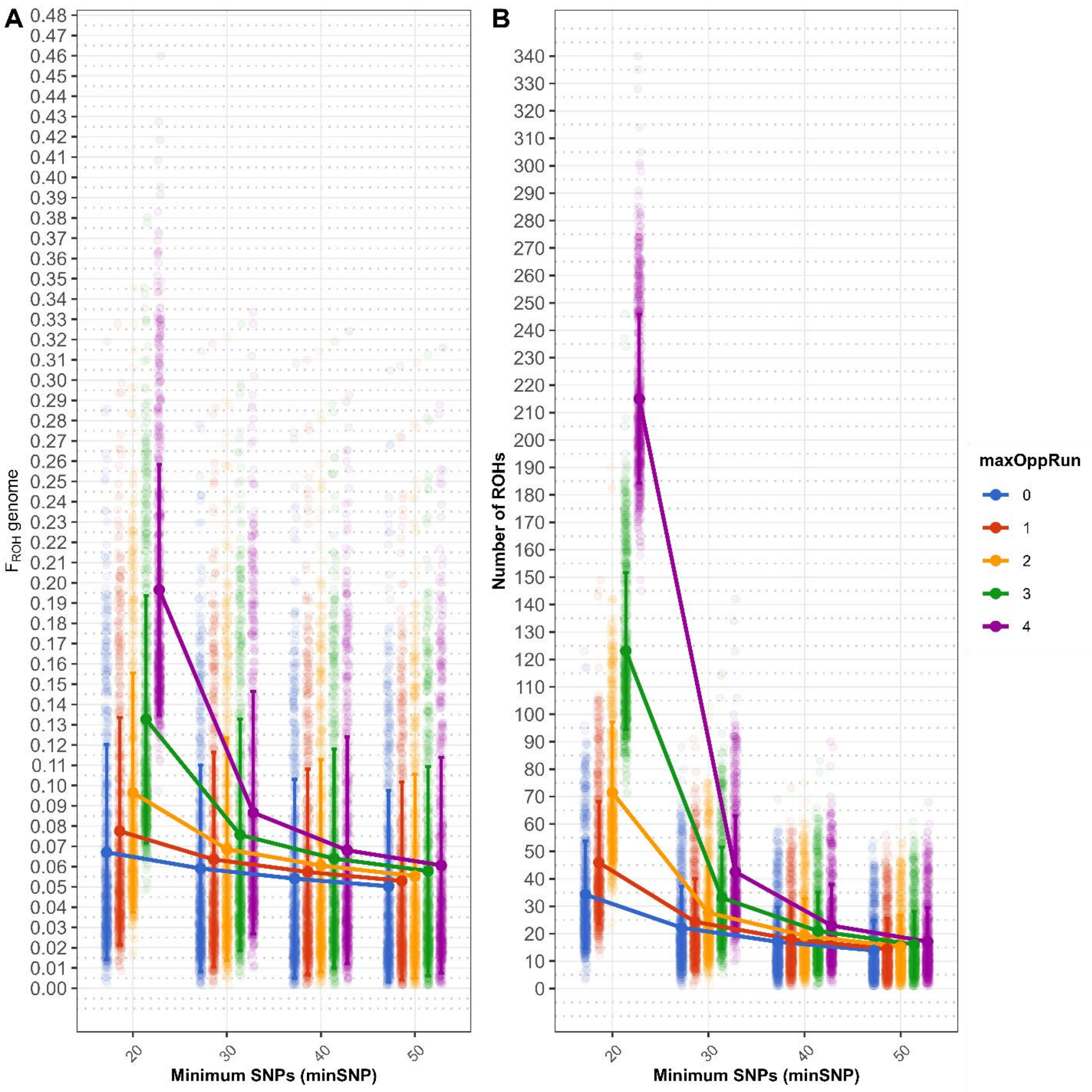


*Figure S1. Effects of parameter settings on ROHs detection in caribou.* (A) Mean $F_{ROH}$ genome values (with standard deviation error bars) across different minimum SNP thresholds and maximum number of heterozygous SNPs (*maxOppRun*). Each colored point represents an individual caribou sample (alpha = 0.05), with solid lines connecting mean values for each parameter combination. (B) Mean number of ROH runs detected per individual under the same parameter combinations. Both panels show results with *maxMissRun* fixed at 1.

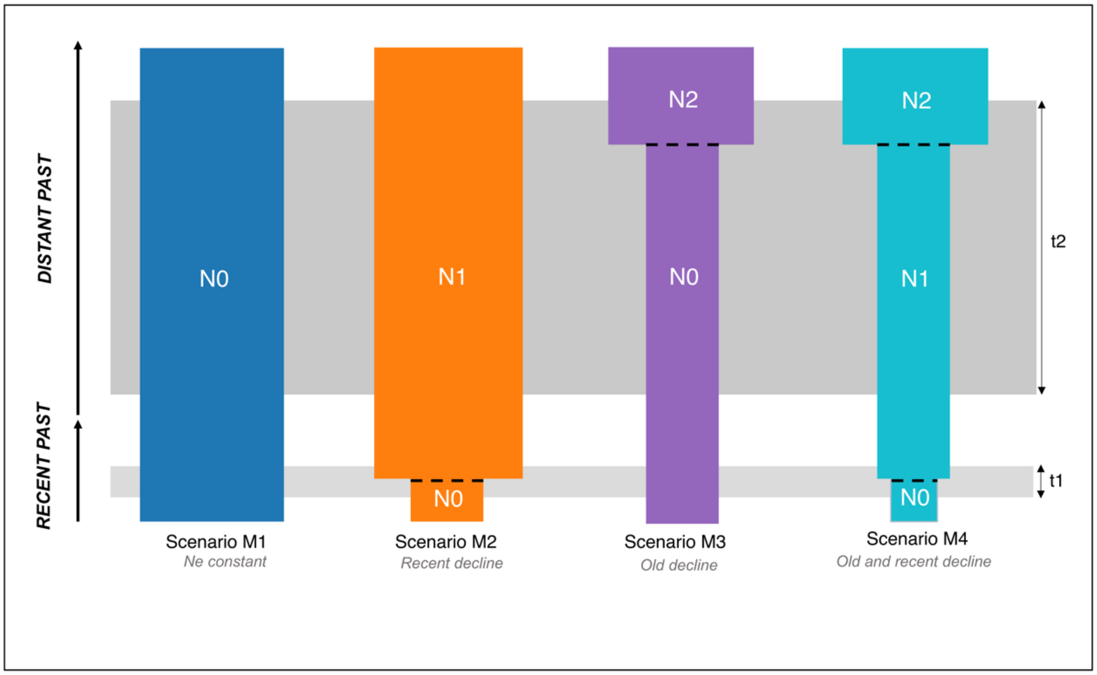


***Figure S2: Bottleneck demographic scenarios tested for 32 caribou subpopulations with DIYABC-RF.*** Only subpopulations with n > 5 were used for this analysis. See Materials & Methods for description of demographic (Ne, N1 and N2) and temporal (t1 and t2) parameters.

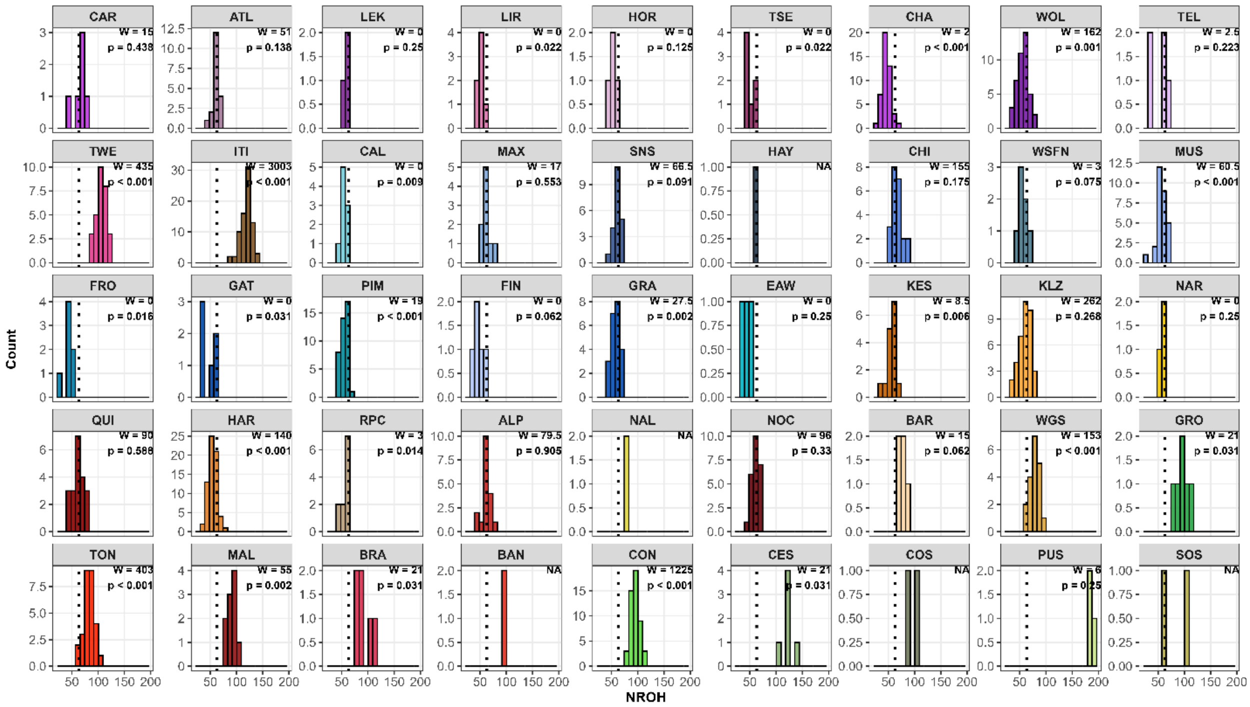


***Figure S3: Description of the number of runs of homozygosity ($N_{ROH}$) detected into 759 individuals across 45 caribou subpopulations in western Canada.*** One-sample Wilcoxon signed-rank tests were performed to compare each subpopulation's values against the overall median. The dotted line corresponds to the overall median.

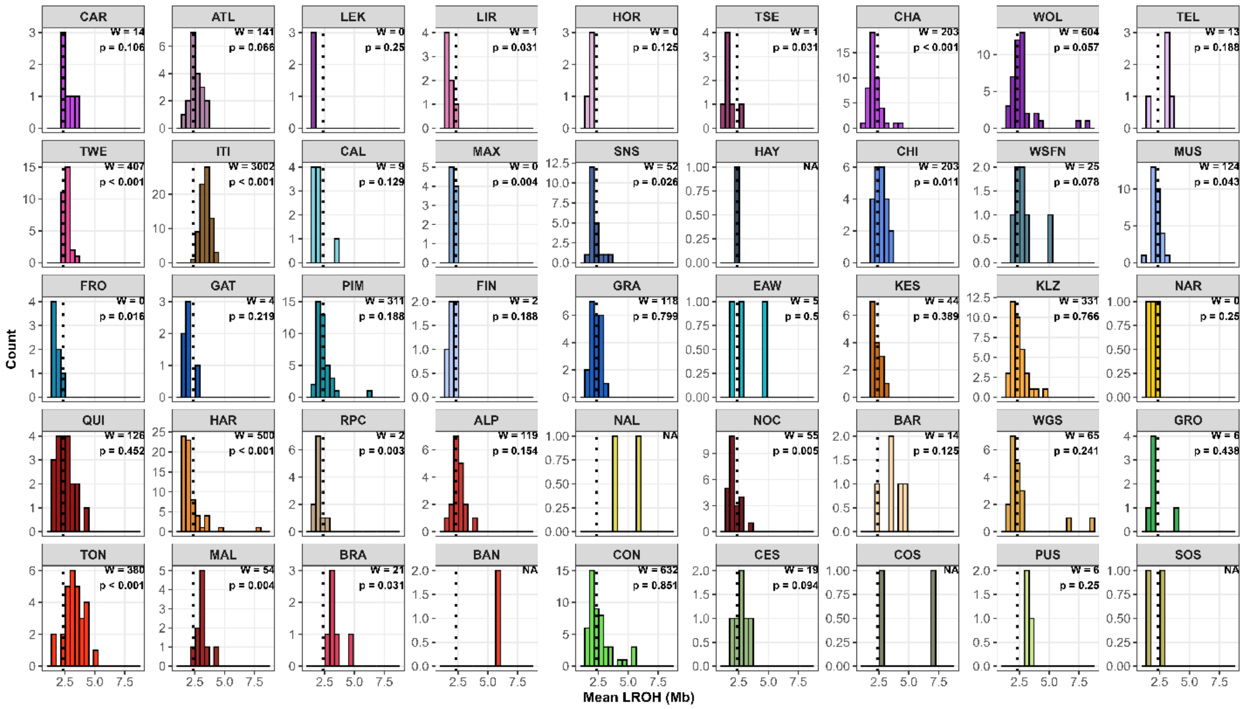


*Figure S4: Distribution of individual mean length of runs of homozygosity ($L_{ROH}$) detected in 759 individuals across 45 caribou subpopulations in western Canada.* One-sample Wilcoxon signed-rank tests were performed to compare each subpopulation's values against the overall median. The dotted line corresponds to the overall median.

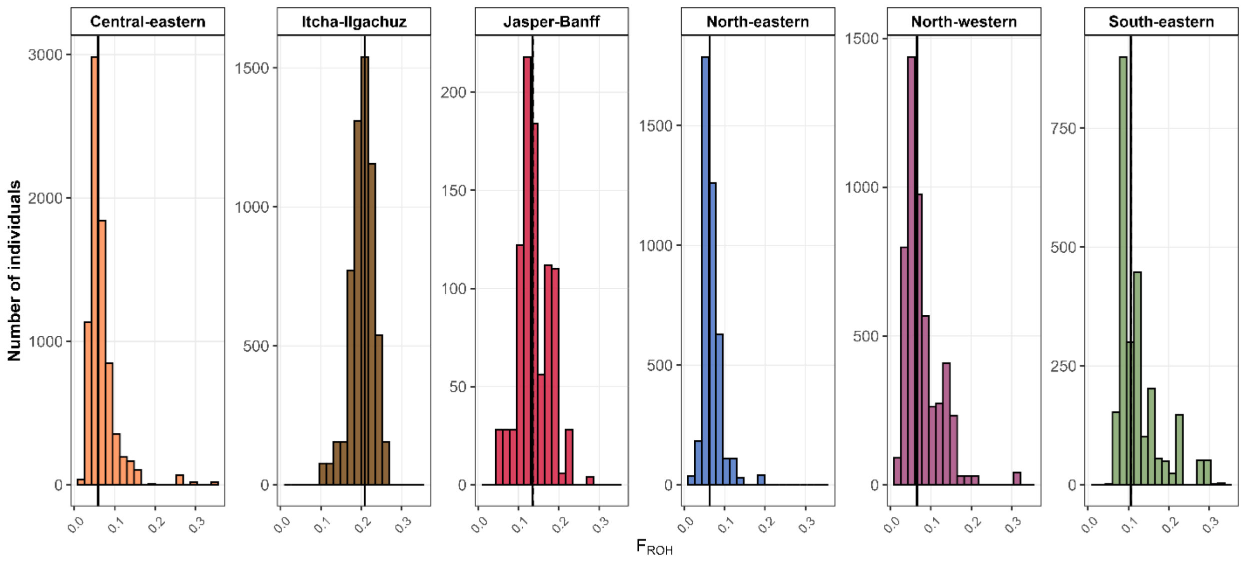


*Figure S5: Distributions of inbreeding coefficients $F_{ROH}$ across 6 metapopulations of caribou in western Canada.* The metapopulations were identified in Deakin et al. (2025). The vertical line corresponds to the median, and the dotted line to the SD.

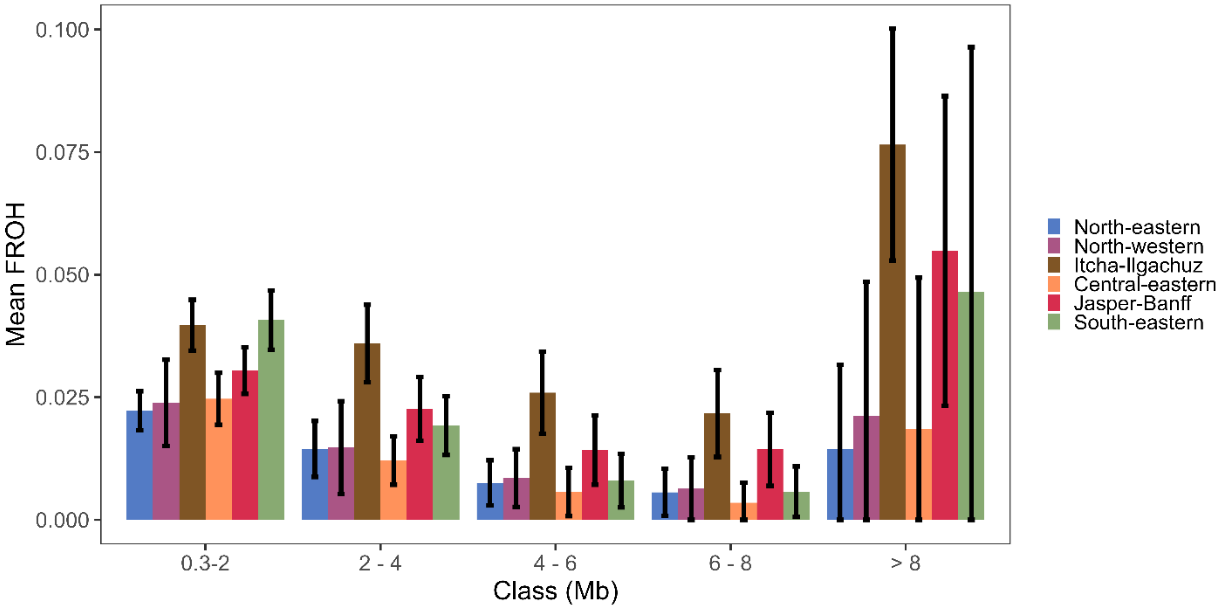


*Figure S6: Inbreeding coefficient derived from Runs of Homozygosity ($F_{ROH}$) per class of ROH lengths from 6 metapopulations of caribou in western Canada.* The different classes were defined as: short (300 kb - 2Mb), medium (between 2-4 Mb, 4-6 Mb and 6-8 Mb) and long (> 8 Mb). Black lines denote standard errors.

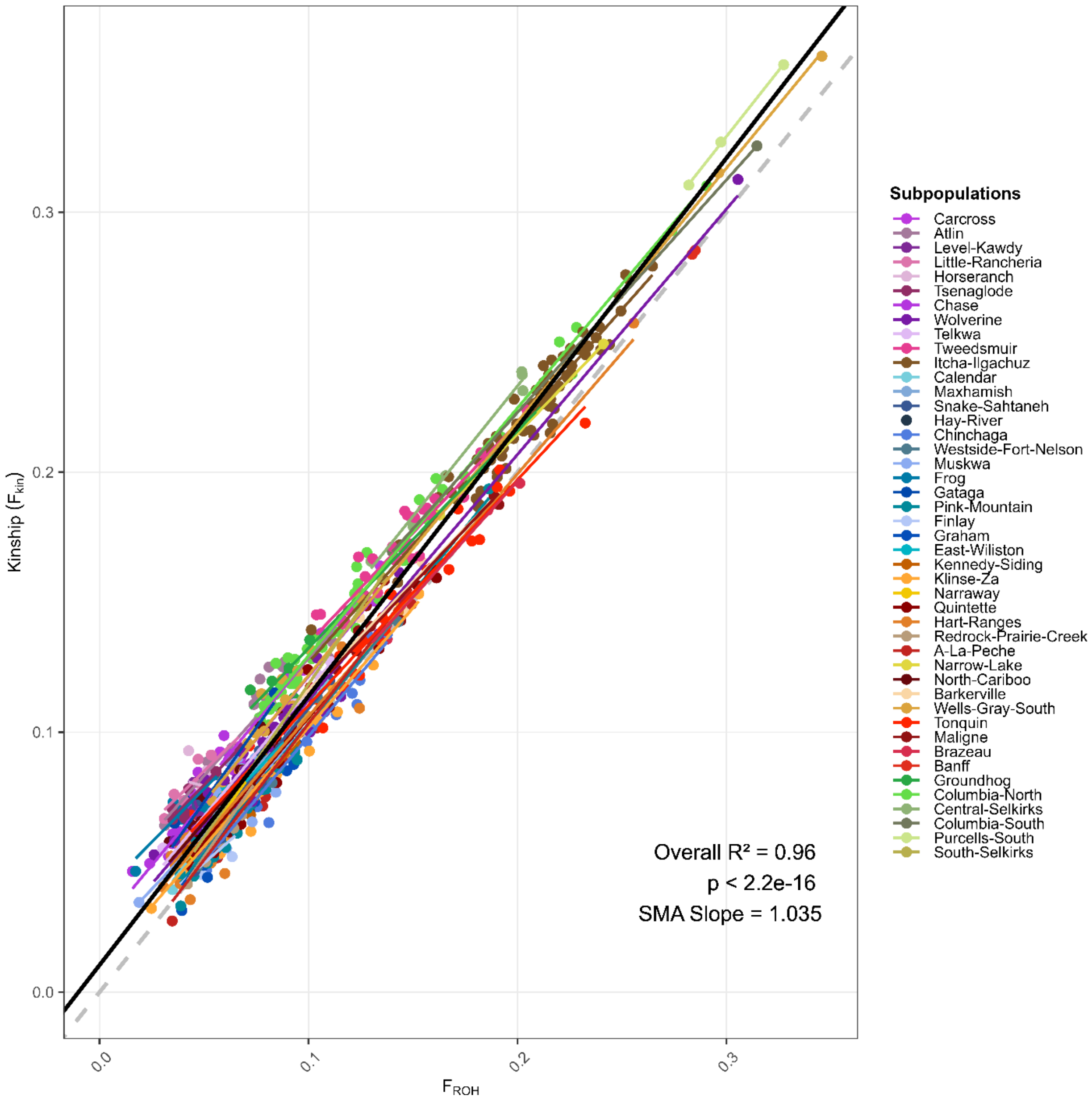


*Figure S7: Relationship between genomic $F_{ROH}$ and $F_{kin}$ across caribou subpopulations.* Each point represents an individual, colored by subpopulation. The dashed gray line represents the 1:1 relationship, while subpopulation-specific linear regressions (colored lines). The black line indicates the overall linear regression with a shaded confidence interval. The inset text provides the overall model fit ($R^2$), statistical significance, and regression slope.

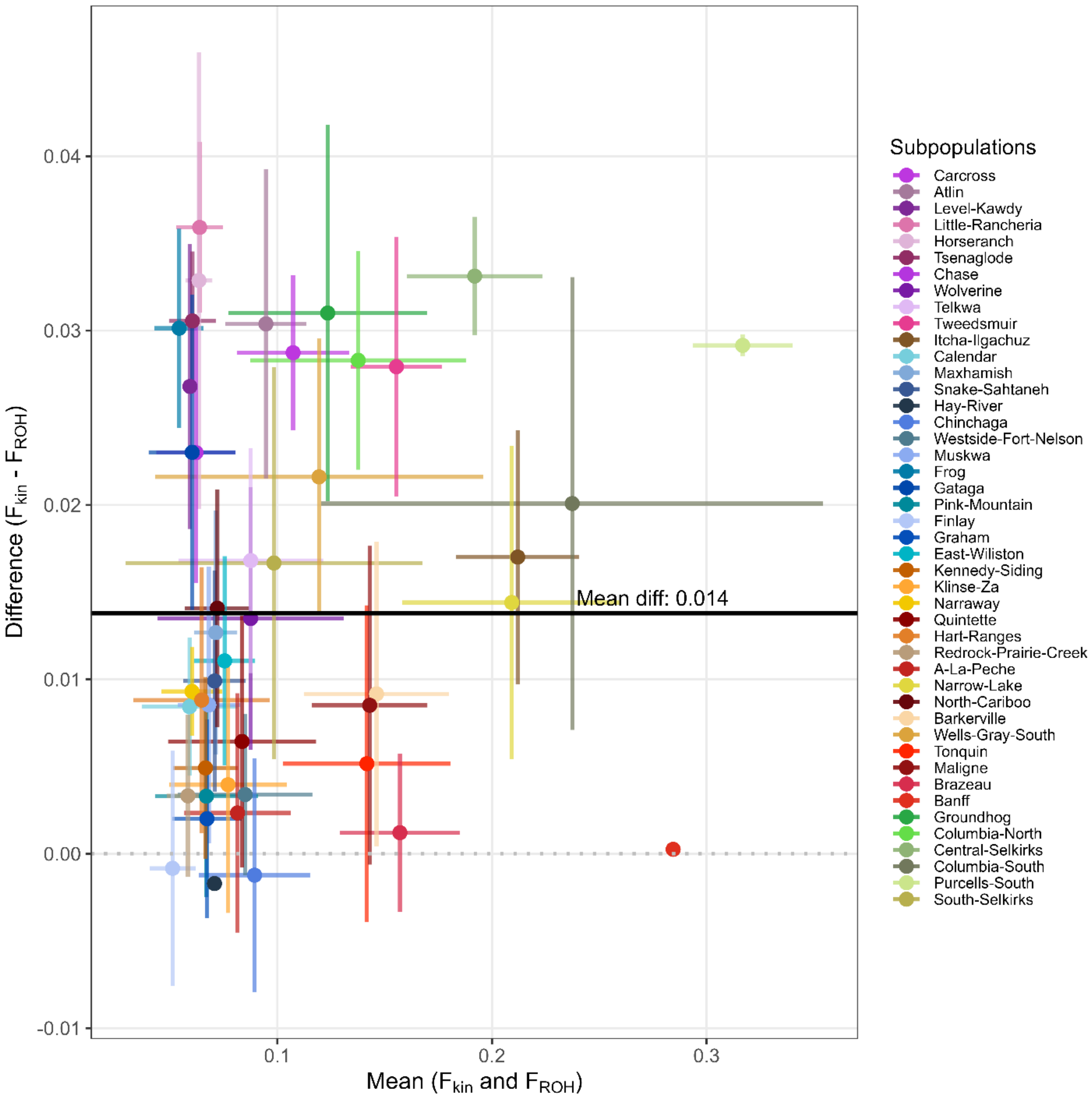


*Figure S8: Bland-Altman plot comparing genomic $F_{ROH}$ and $F_{kin}$ across caribou subpopulations.* Each point represents the mean difference between $F_{ROH}$ and $F_{kin}$ for a subpopulation, with horizontal and vertical error bars indicating one standard deviation of the mean and difference, respectively. The solid black line represents the mean difference, while the dotted gray line marks zero difference.

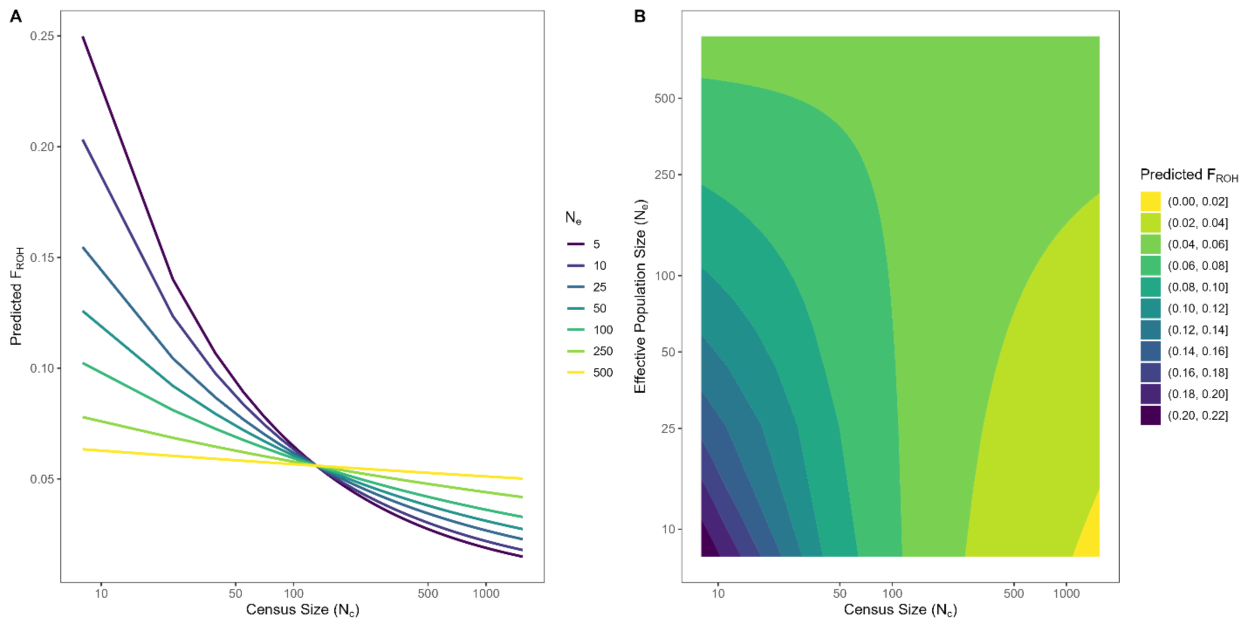


*Figure S9: Interaction effect of census size ($N_c$) and effective population size ($N_e$) on inbreeding coefficient ($F_{ROH}$) in caribou subpopulations.* A. Predicted relationship between $N_c$ and $F_{ROH}$ at different levels of $N_e$. Lines represent predicted $F_{ROH}$ values across a range of $N_c$ values while holding $N_e$ constant at different levels (5-500). B. Contour plot visualizing the combined effect of $N_c$ and $N_e$ on predicted mean $F_{ROH}$. Color gradient represents predicted $F_{ROH}$ values, with lighter yellow areas indicating lower inbreeding and darker blue/purple areas indicating higher inbreeding.

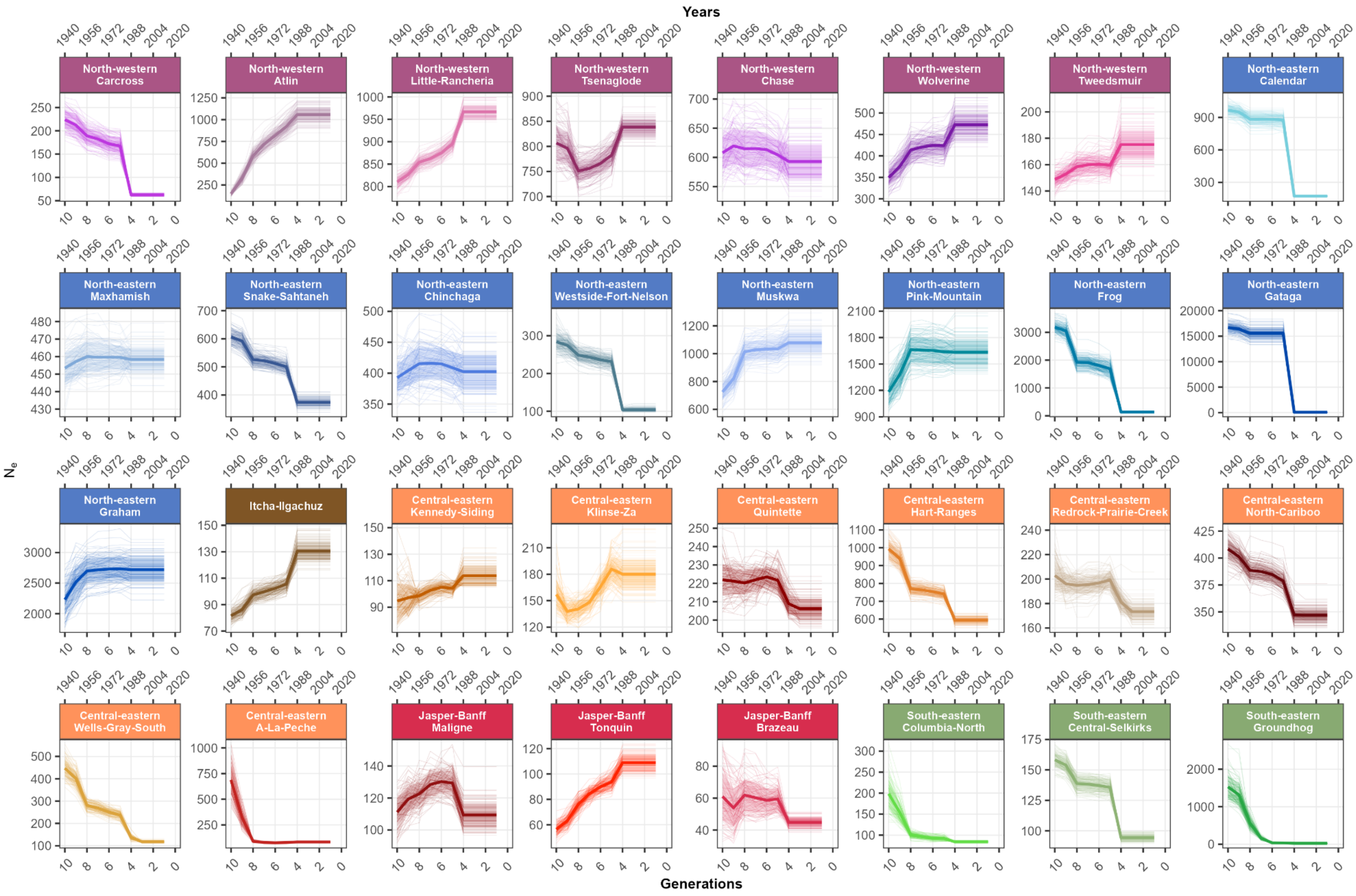
Years
$N_e$
Generations
North-western Carcross
North-western Atlin
North-western Little-Rancheria
North-western Tsenaglode
North-western Chase
North-western Wolverine
North-western Tweedsmuir
North-eastern Calendar
North-eastern Maxhamish
North-eastern Snake-Sahtaneh
North-eastern Chinchaga
North-eastern Westside-Fort-Nelson
North-eastern Muskwa
North-eastern Pink-Mountain
North-eastern Frog
North-eastern Gataga
North-eastern Graham
Itcha-Ilgachuz
Central-eastern Kennedy-Siding
Central-eastern Klinse-Za
Central-eastern Quintette
Central-eastern Hart-Ranges
Central-eastern Redrock-Prairie-Creek
Central-eastern North-Cariboo
Central-eastern Wells-Gray-South
Central-eastern A-La-Peche
Jasper-Banff Maligne
Jasper-Banff Tonquin
Jasper-Banff Brazeau
South-eastern Columbia-North
South-eastern Central-Selkirks
South-eastern Groundhog

***Figure S10: Historical effective population size ($N_e$) trajectories over the last 10 generations (~80 years) for 32 caribou (Rangifer tarandus) subpopulations in western Canada.*** $N_e$ point estimates were obtained using GONE2 v2.0 under a panmictic model. The thick solid lines represent the mean $N_e$ estimates calculated from 100 independent stochastic runs (seeds) at a recombination rate of 1.0 cM/Mb. Shaded ribbons represent ± 1 standard deviation around the mean. Subpopulations are grouped and colored by their corresponding metapopulation as defined by Deakin et al. (2025) based on genetic structure analyses. Subpopulations with small sample sizes ($n \leq 5$) were excluded from analysis. Generations were converted to years (upper x-axis), assuming a mean generation time of 8 years.

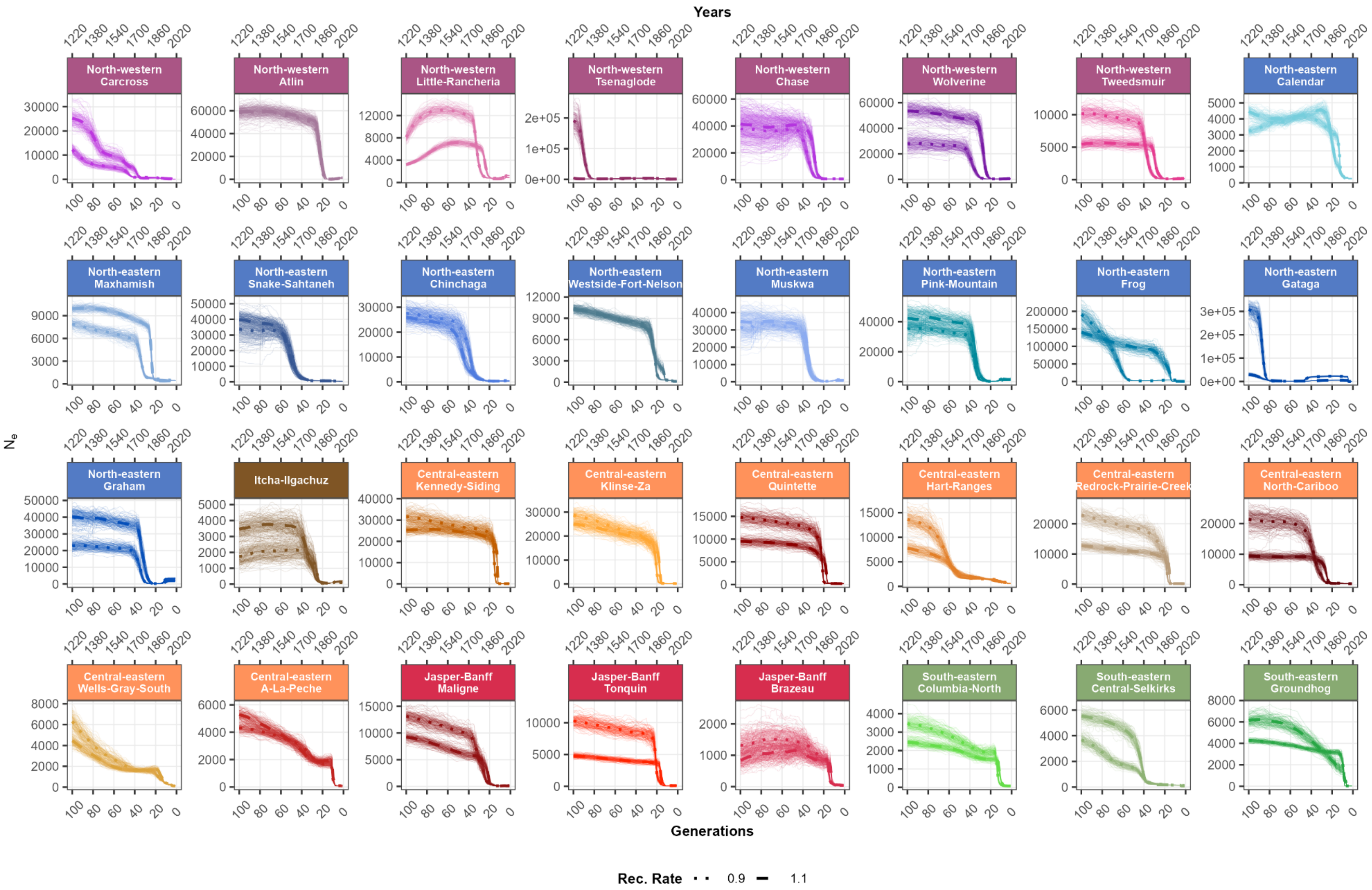
Years
North-western Carcross
North-western Atlin
North-western Little-Rancheria
North-western Tsenaglode
North-western Chase
North-western Wolverine
North-western Tweedsmuir
North-eastern Calendar
North-eastern Maxhamish
North-eastern Snake-Sahtaneh
North-eastern Chinchaga
North-eastern Westside-Fort-Nelson
North-eastern Muskwa
North-eastern Pink-Mountain
North-eastern Frog
North-eastern Gataga
North-eastern Graham
Itcha-Ilgachuz
Central-eastern Kennedy-Siding
Central-eastern Klinse-Za
Central-eastern Quintette
Central-eastern Hart-Ranges
Central-eastern Redrock-Prairie-Creek
Central-eastern North-Cariboo
Central-eastern Wells-Gray-South
Central-eastern A-La-Peche
Jasper-Banff Maligne
Jasper-Banff Tonquin
Jasper-Banff Brazeau
South-eastern Columbia-North
South-eastern Central-Selkirks
South-eastern Groundhog
$N_e$
Generations
Rec. Rate 0.9 1.1

***Figure S11: Historical effective population size ($N_e$) trajectories over the last 100 generations (~800 years) for 32 caribou (Rangifer tarandus) subpopulations in western Canada, comparing different recombination rates.*** *$N_e$ point estimates were obtained using GONE2 v2.0 under a panmictic model.* Solid lines represent the mean $N_e$ estimates at a recombination rate of 1.0 cM/Mb, dotted lines at 0.9 cM/Mb, and dashed lines at 1.1 cM/Mb. All estimates were calculated from 100 independent stochastic runs (seeds). Subpopulations are grouped and colored by their corresponding metapopulation as defined by Deakin et al. (2025) based on genetic structure analyses. Subpopulations with small sample sizes ($n \leq 5$) were excluded from analysis. Generations were converted to years (upper x-axis), assuming a mean generation time of 8 years.

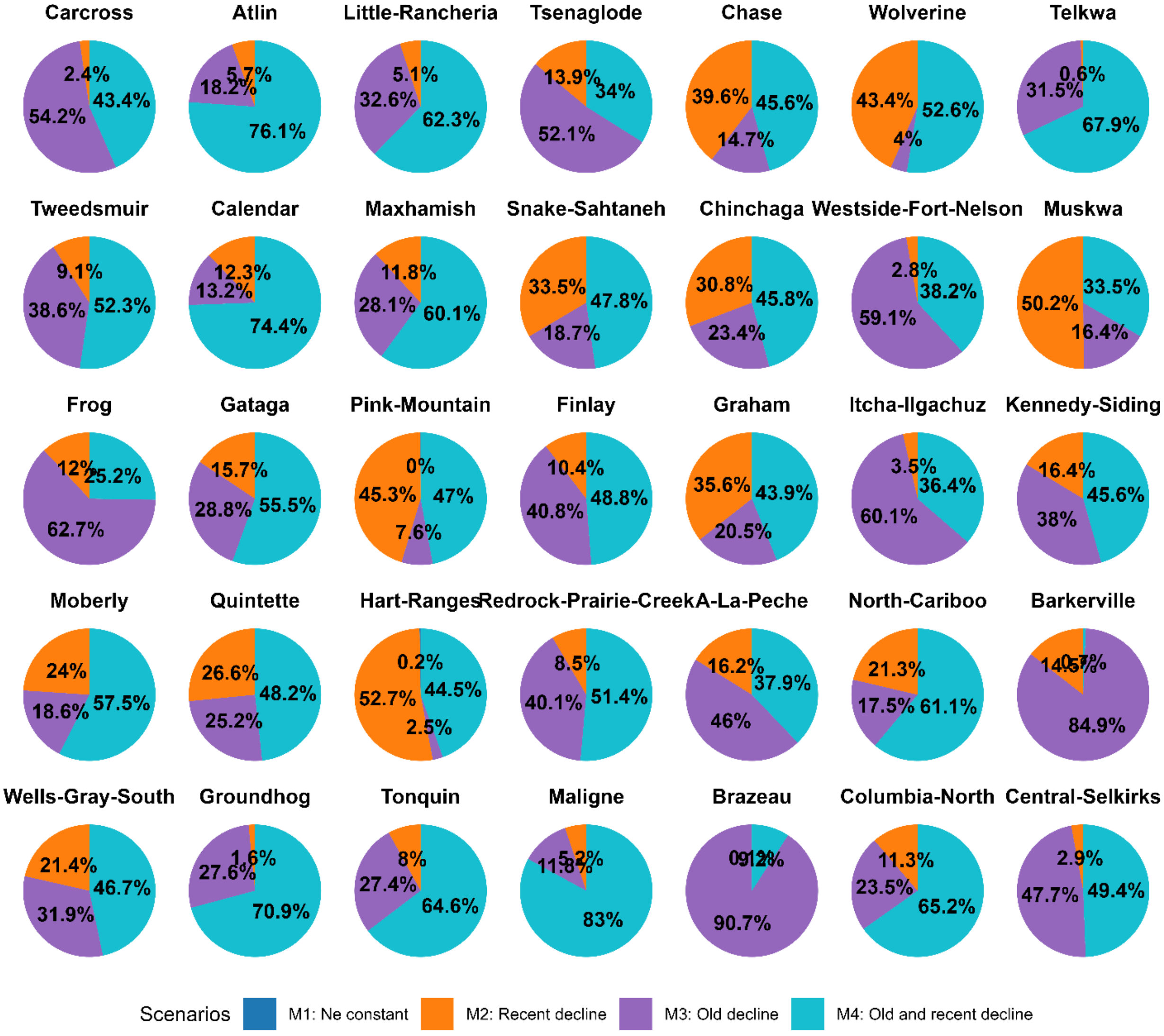


*Figure S12: Demographic scenario votes across 32 caribou subpopulations.* Each pie chart represents the proportion of 2000 votes assigned to the competing scenarios: $N_e$ constant (Scenario M1; dark blue), Recent decline (Scenario M2, orange), Old decline (Scenario M3, purple), and Old and recent decline (Scenario M4, turquoise). Analyses were conducted for 32 subpopulations with $n > 5$.
